# Ritornello: High fidelity control-free chip-seq peak calling

**DOI:** 10.1101/034090

**Authors:** Kelly Patrick Stanton, Jiaqi Jin, Sherman Weissman, Yuval Kluger

## Abstract

With the advent of next generation high-throughput DNA sequencing technologies, omics experiments have become the mainstay for studying diverse biological effects on a genome wide scale. ChIP-seq is the omics technique that enables genome wide localization of transcription factor binding or epigenetic modification events. Since the inception of ChIP-seq in 2007, many methods have been developed to infer ChIP target binding loci from the resultant reads after mapping them to a reference genome. However, interpreting these data has proven challenging, and as such these algorithms have several shortcomings, including susceptibility to false positives due to artifactual peaks, poor localization of binding sites, and the requirement for a total DNA input control which increases the cost of performing these experiments. We present Ritornello, a new approach with roots in digital signal processing (DSP) that addresses all of these problems. We show that Ritornello generally performs equally or better than the peak callers tested and recommended by the ENCODE consortium, but in contrast, Ritornello does not require a matched total DNA input control to avoid false positives, effectively decreasing the sequencing cost to perform ChIP-seq.

## Introduction

Reliable and precise characterization of where proteins, such as transcription factors interact with the genome, enables biologists to understand how gene expression is regulated at the molecular level. The human genome, for example, encodes about 1500 transcription factors (TFs) [1] and many of them directly recognize and bind to specific DNA sequences to regulate gene expression. Therefore, identification of where each TF binds to the DNA is critical for reconstructing the complex regulatory network of gene expression. Chromatin immunoprecipitation (ChIP) followed by high-throughput sequencing (ChIP-seq) is a powerful tool for detecting protein-DNA interactions at the genome-wide scale and has become the method of choice. In a ChIP-seq experiment, first, proteins interacting with the DNA are chemically attached to the DNA using formaldehyde-mediated crosslinking. Then the DNA is fragmented into short pieces and antibodies specifically targeting the protein of interest are used to pull down DNA fragments bound by that protein. Finally, the immunoprecipitated DNA fragments are released from the protein of interest and subjected to high-throughput DNA sequencing. The resulting sequenced reads are mapped to a reference genome and computational algorithms are applied to process mapped reads and infer protein binding positions (peak calling).

Transcription factors usually bind to short specific DNA sequences (motifs) and generate sharp point-source peaks [2]. For most ChIP-seq experiments currently available, only one of the two 5’ ends of each doublestranded DNA fragment has been sequenced (single end sequencing), so the read coverage near the point-source peaks follow a characteristic bimodal shape. However, calling peaks accurately from a large quantity of mapped reads is nontrivial and over 40 algorithms have been developed [3–44] since the ChIP-seq technology was first introduced [45]. Peak calling remains challenging due to the presence of artifactual binding events (false positives) and background noise from reads outside of peaks, multi-binding events with overlapping read contributions, and variability of experimental quality. Additionally, for most peak calling algorithms, matched negative controls, which are usually DNA samples obtained without performing immunoprecipitation (total DNA input control) or immunoprecipitated by non-specific antibodies (IgG control), are often required to control the false positive rate.

Performing a negative control experiment for each sample, effectively doubles the sequencing cost of ChIP-seq, limiting the number of samples that can be run per experiment. Peak calling algorithms that do not use the control (including those that have the option to run with or without it) have been developed, however, they underperform due to the lack of a detailed characterization of ChIP-seq signal and noise.

Binding events can also occur in close proximity to one another, and it is often difficult to resolve how many binding sites are present and precisely where binding occurs. BRACIL [46] and CSDeconv [47] use blind deconvolution algorithms to resolve individual peaks at multi-binding loci but are not scalable for peak calling and are thus used for post processing when peaks have been identified by other peak callers. GEM [23] incorporates *de novo* motif discovery into the peak identification process aiding in resolving individual peaks, but may not be suitable if the transcription factor of interest does not bind to DNA directly or does not have any specific motif.

ChIP-seq experiments can also be of varying quality. Collective efforts by large consortia have provided guidelines on how to evaluate the quality and signal-to-noise ratio of ChIP-seq experiments. The opposing strand cross-correlation between the read coverage on the positive and that on the negative strands has been used to assess experimental quality by ENCODE [2]. The cross-correlations of ChIP-seq as well as input control experiments exhibit two modes, one at or around their respective average fragment lengths, and an additional one at or around their respective read lengths. High quality experiments tend to have a greater contribution from the fragment length mode, while low quality experiments and input controls tend to have a larger contribution from the read length mode. Specifically, the ENCODE ChIP-seq guidelines include two metrics: the normalized strand coefficient(NSC) and the relative strand correlation(RCS) [2]. If the NSC or RSC scores are low, indicating poor experimental quality, ENCODE recommends repeating the experiment. Given the considerable cost of repeating a ChIP-seq experiment, it is useful to be able to “rescue” samples with suboptimal quality for use as additional replicates, rather than discarding them.

Here, we present Ritornello, a novel ChIP-seq peak calling algorithm based on both digital signal processing and statistical techniques. In the current work, we contribute the following innovations and insights:

- a peak caller, that does not require a matched control and still maintains a low false positive rate, outperforming even algorithms that use the control
- an efficient method to perform full deconvolution of multi-binding events on a genome wide scale
- samples of low quality can be “rescued”, instead of being discarded
- a rigorous characterization of the binding signals and artifacts in the presence of noise in ChIP-seq data
- a nonparametric approach to calculate the fragment length distribution for any single-end NGS experiment.

We benchmarked Ritornello against MACS2 [3] and GEM [23], two algorithms recommended by the ENCODE consortium [2]. In the default modes each requires the matched control. We demonstrated that Ritornello, a matched control free method, outperformed MACS2 and GEM.

## Methods

We have developed Ritornello to find candidate peaks efficiently, with minimal use of memory and computation time, by using a digital signal processing technique called a matched filter, classify candidate peaks as true binding events or artifacts based on their shape, and finally test candidate binding positions for significance based on comparison to a model absent of binding at that position. The scheme of the Ritornello method is detailed in Figure 1.

**Figure 1:**
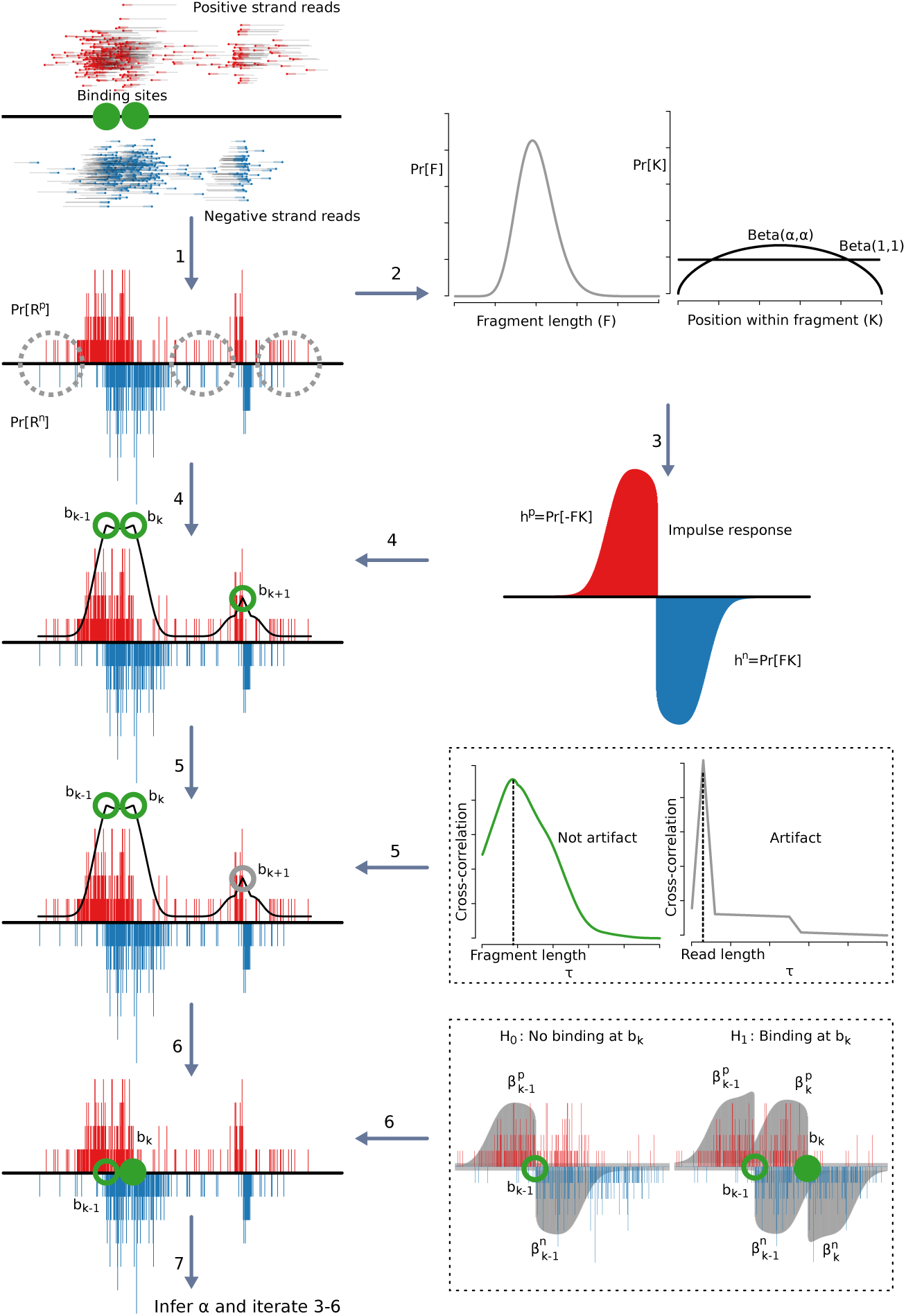
Overview of the Ritornello approach. Step 1: the reads are mapped to the genome and the distributions of read starts Pr[*R^p^*] and Pr[*R^n^*] are calculated. Step 2: the fragment length distribution *Pr*[*F*] is calculated using only those areas where the requisite independence assumptions, *R^p^* ⊥ *F* or *R^n^* ⊥ *F*, hold (background coverage). Step 3: the expected distribution of reads around a binding event is calculated from a model including the fragment length distribution Pr[*F*] as well as the distribution of relative binding positions within fragments Pr[*K*]. Initially, *K* is modeled with a uniform distribution, or equivalently *K* ~ *Beta*(*α, α*) with *α* = 1. Step 4: the expected distribution of reads around a binding event is used to locate candidate binding event peaks (e.g. *b_k−1_, b_k_*, and *b_k_*_+1_) by identifying the positions with the highest match with impulse response function. Step 5: candidate binding events are classified as either read length artifacts (e.g. *b_k+1_*) or are retained as putative binding events (e.g. *b_k−_*_1_ and *b_k_*) based on the shape of the local cross-correlation between opposing strands. Step 6: the binding intensity, 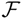, of each peak, *b_k_* are deconvolved from the mixture of local peaks *b_k−_*_1_ and background noise using a maximum likelihood approach. The likelihood ratio test is then applied to determine the significance of the peak. Step 7: the distribution of relative binding positions within fragments, Pr[*K*], is updated using a maximum likelihood estimate of *α*, where *K* ~ *Beta*(*α, α*). This is obtained using a combined likelihood model for the top 200 most significant peaks. Steps 3–6 are repeated using the new Pr[*K*].

### Derive fragment length distribution via deconvolution

For fragments overlapping a binding position, the positive strand mapping reads will be upstream of the binding site whereas negative strand reads will be downstream of the site. As such, the distance between a read and the binding position is dependent on the fragment length, which most peak calling algorithms must estimate (usually mean fragment length) to obtain accurate predictions for binding locations. Our first innovation for Ritornello is calculating, not just the mean fragment length, but the entire sample specific empirical fragment length distribution (FLD) from single-end reads (step 2 of Figure 1). Ritornello utilizes this FLD, as a key component, for more accurate peak predictions, which we will describe in detail below.

The cross-correlation is the single strand autocorrelation convolved against the fragment length distribution as has previously been shown [48]:

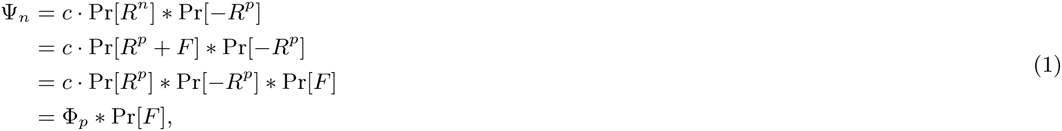
where Ψ*n* is the opposing strand cross-correlation, Φ*_p_* is the autocorrelation of the positive strand, Pr[*R^p^*] is the probability of choosing a read starting at a position *R^p^* on the positive strand, Pr[*R^n^*] is the probability of choosing a read start at a position *R^n^* on the negative strand, Pr[*F*] is the probability of sampling a fragment of length *F*, (fragment length distribution), *c* is the read count, and * is the convolution operator. The fragment length distribution can then be obtained by deconvolution as follows:

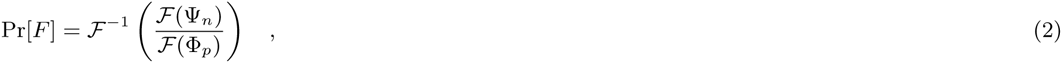
where 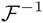 is the Fourier transform operator and 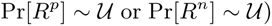 is its inverse.

In the current work, we clarify that **Equation** 1 assumes that *R^p^* and *F* are statistically independent, and thus Pr[*R^p^* + *F*] can be simplified to Pr[*R^p^*] * Pr[*F*]. If we assume that *R^n^* (as opposed to *R^p^*) and *F* are independent, then we can instead write the following relationship:

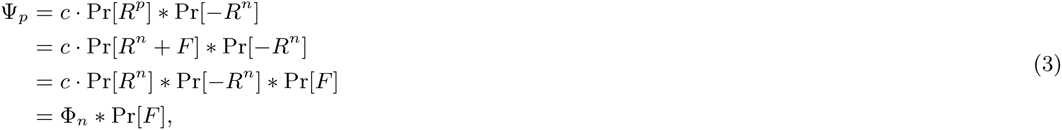
and deconvolve as follows:

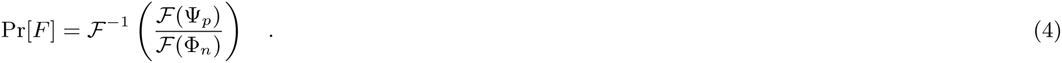

**Equations** 2 and 4 describe how to obtain the fragment length distribution under two different and mutually exclusive assumptions (i.e. *R^p^* ⊥ *F* or *R^n^* ⊥ *F* where ⊥ denotes statistical independence). To estimate the fragment length distribution, Ritornello locates genomic positions where either *R^p^* ⊥ *F* or *R^n^* ⊥ *F* and invokes the corresponding equation locally. However, it is highly likely that regions, outside of binding events, with relatively uniformly distributed reads on *R^p^* or *R^n^* satisfy *R^p^* ⊥ *F* or *R^n^* ⊥ *F* respectively.

We identify these regions by looking for read coverage that is locally uniform on either strand, using a χ^2^ goodness of fit test (i.e. 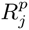). For each window of size *2F_max_* (twice the maximum fragment length) centered at position *i* on either strand, we calculate the χ^2^ test statistic as follows:

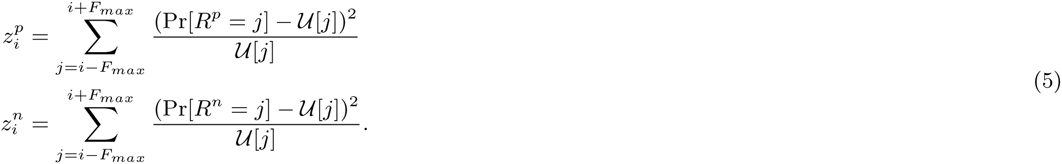

We then sum the local autocorrelations, Φ*_p,i_* or Φ*_n,i_*, for those windows where either the positive or negative strand is independent of the fragment length, as determined by the *χ*^2^ test for local uniformity, across the genome *G*. Additionally, we sum the local opposing strand cross-correlations, Ψ*_p,i_* or Ψ*_n,i_*, associated with each autocorrelation according to **Equations** 1 and 3 as follows:

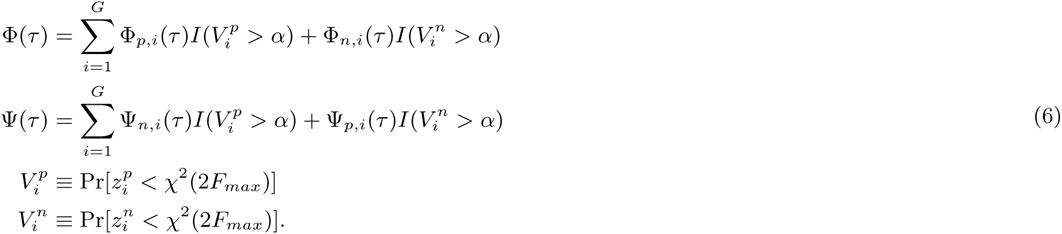
where,

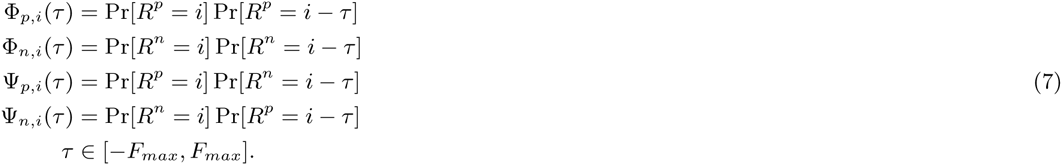

Using the global autocorrelation, Φ, and cross-correlation, Ψ, functions from **Equation** 6, we calculate the fragment length distribution as follows:

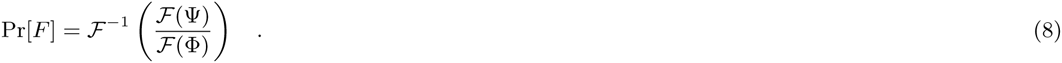

### Why the fragment length distribution can be inferred from single end data?

We have just shown that the fragment length distribution can be derived via deconvolution from Pr[*R^p^*] and Pr[*R^n^*] when those distributions are related by **Equations** 1 or 3. This is intuitive in paired-end sequencing. However, in single-end sequencing, only one end is randomly selected from each fragment, and it is hard to imagine how the fragment length information can be preserved. For a given set of fragments, single end sequenced reads are a sub-sample of paired-end sequenced reads. Thus, Pr[*R^p^*] and Pr[*R^n^*], the only input to **Equations** 2 or 4, in the single-end reads should be the same with that in the paired-end, resulting in the same estimated fragment length distribution, Pr[*F*]. A reliable approximation of the paired end distributions Pr[*R^p^*] and Pr[*R^n^*] are obtained from locations where multiple fragments share either a common start or end position (within a couple of base pairs) such as in Figure 2a. Collectively, all such locations in the genome enable a faithful reconstruction of the whole fragment length distribution as in paired-end sequencing (Figure 2a).

**Figure 2:**
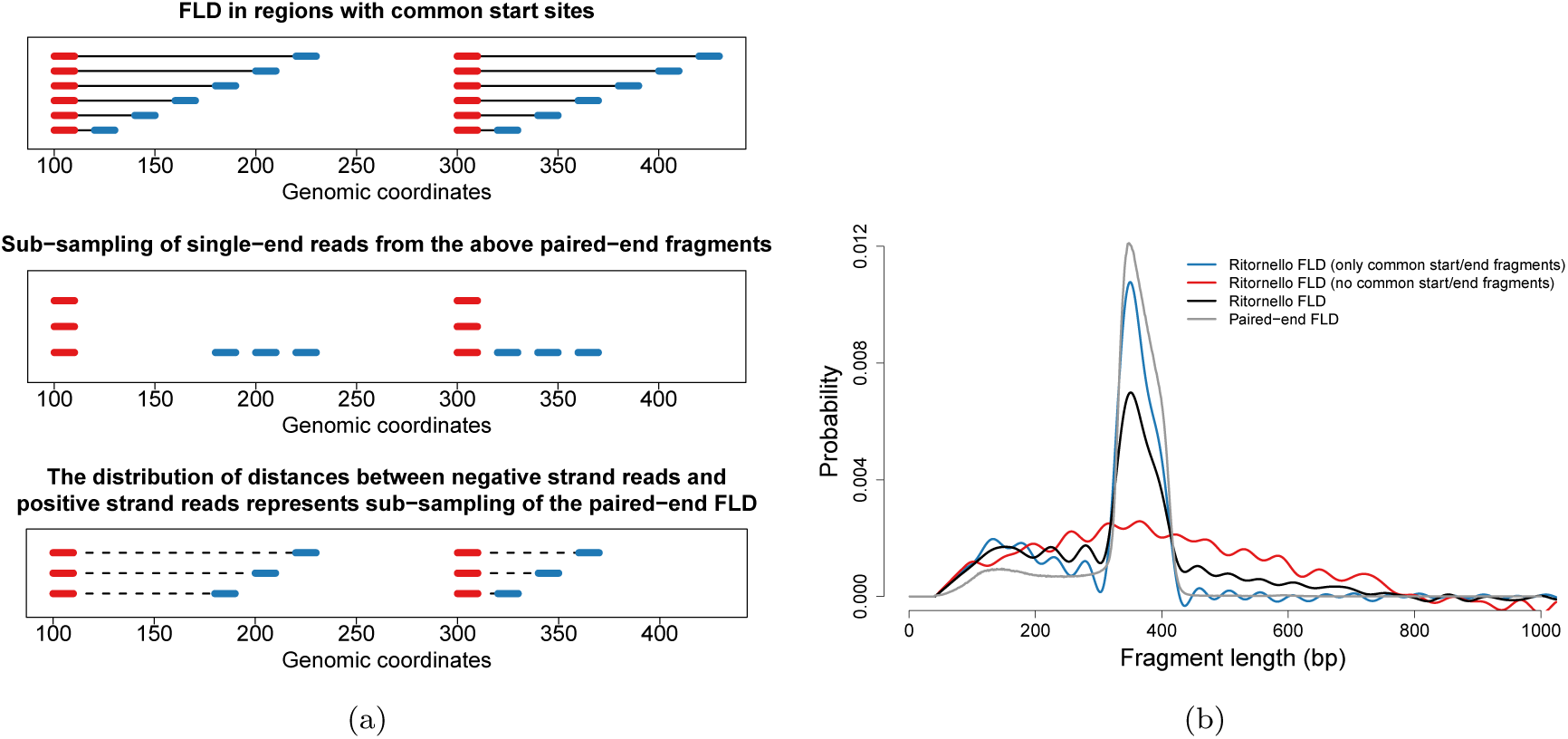
Ritornello captures the fragment length distribution from single-end sequencing data. Given a set of fragments, singled end sequenced reads are a subsample of paired-end sequenced reads. When enough single end fragments are sampled the distribution of read coverage on the positive and negative strands Pr[*R^p^*] and Pr[*R^n^*] are equivalent to their paired-end counterparts. For ChIP-seq, outside of binding events and artifacts, this occurs most often in areas where fragments share common start or end positions. (a) an illustrative example of reconstruction of the fragment length distribution using fragments sharing a common start position. Paired-end fragments are simulated to form a common start position (panel 1 in subfigure a). Single end reads are subsampled from the same fragments used for paired end (panel 2 of subfigure a). The distances between opposing strand single end reads originating from a set of fragments that share a common start position mimic the fragment lengths (b) Accurate fragment length distribution calculation from single end data depends on fragments with common start or end positions. The fragment length distribution calculated by Ritornello from a paired-end EZH2 sample subsampled to simulated singled end data is shown in black. The fragment length structure is lost when reads belonging to fragments that share common start or end positions are discarded (red curve). The fragment length distribution calculated from the subset of fragments that share a common start (blue curve), closely approximates the true fragment length distribution (gray curve).

To demonstrate this point we have isolated fragments sharing common start or end positions in a paired-end total DNA input, randomly sampled one end from each fragment to create a pseudo single-end data, and used the deconvolution approach to calculate fragment length distribution. In Figure 2b, we show that the fragment length distribution calculated from this pseudo single-end data (blue) is very similar to the true fragment length distribution of the paired-end sample (green). When pseudo single-end data is created from all fragments in the paired-end data (not only from fragments that share common start or end positions), the fragment length distribution calculated via deconvolution (black) is less similar to the true fragment length distribution. When pseudo single-end data is created from all fragments except those that share common start or end positions, the fragment length distribution calculated via deconvolution (red) dramatically deviates from the true fragment length distribution. This suggests that even in single-end sequencing, the distribution of reads originating from fragments that share common start or end positions reliably approximate the paired end Pr[*R^p^*] and Pr[*R^n^*] distributions, enabling accurate calculation of the fragment length distribution.

### Local fragment length distribution varies around binding events

The fragments generated from a binding event overlap the binding site, thus any given fragment must be at least as long as the distance from its start position to the binding site. This creates dependence between the fragment length and genomic position on both strands because reads that are further from the binding site are necessarily longer on aggregate. Consequently, neither the positive nor the negative strand read coverage is independent of the fragment length within a binding event, making it inappropriate to recover the fragment length distribution using **Equation** 2, as shown by simulation in Figure 3a. In contrast, in simulated event free regions, it is shown that the fragment length distribution can be correctly recovered using **Equation** 2 as seen in Figure 3b.

**Figure 3:**
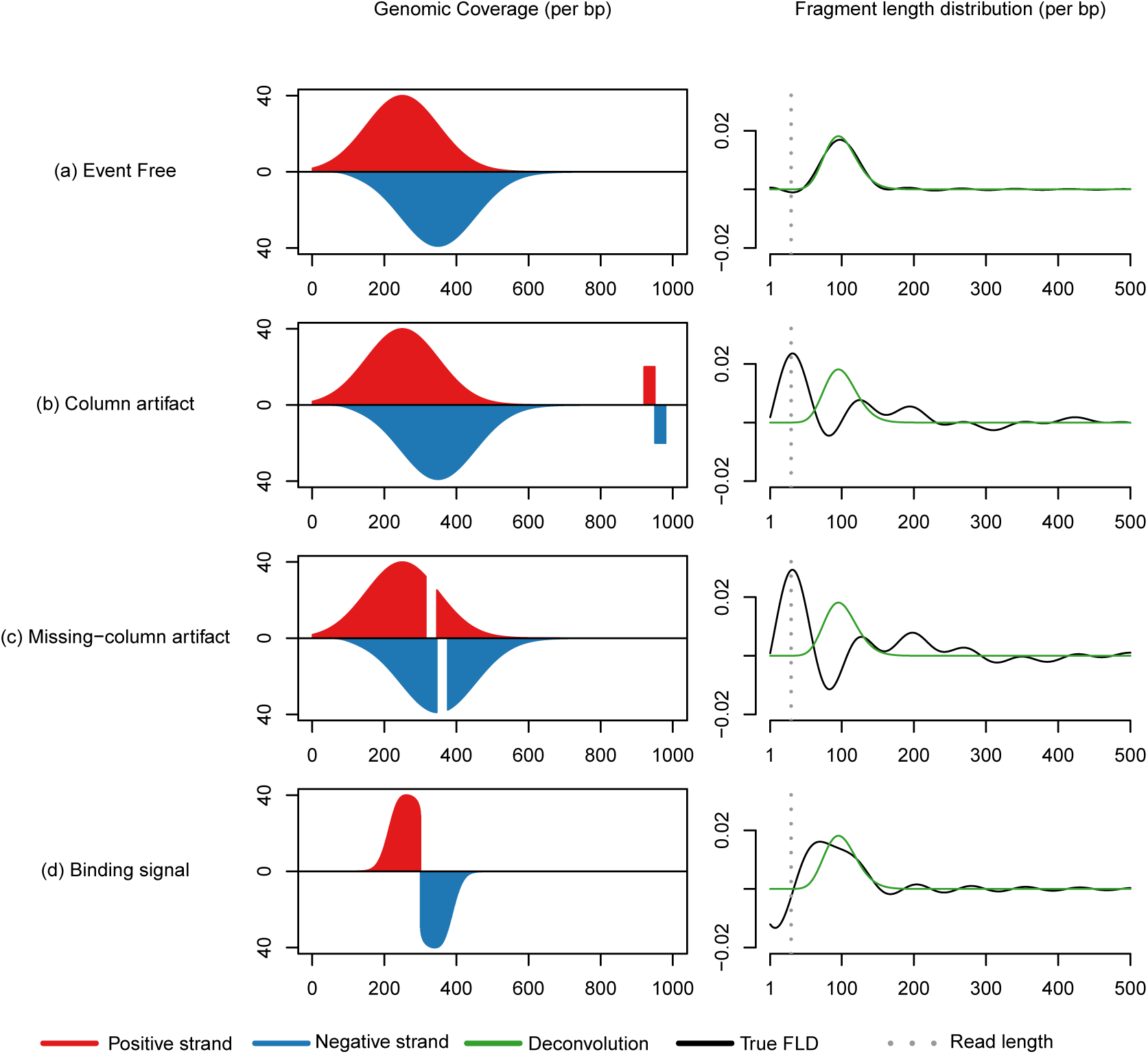
The presence of binding events and read length artifacts hinders reconstruction of the fragment length distribution by deconvolution. Coverage patterns (positive strand in red, negative strand in blue) were generated from reads sampled randomly from either end of simulated fragments. **Equation** 2 was applied to infer the fragment length distribution, FLD (black). The actual FLD was calculated from the simulated fragments (green). The read length is shown in gray. (a) The FLD, inferred from reads in simulated binding regions, deviates from the true FLD. (b) The FLD, inferred from genomic background coverage outside of binding events, agrees with the true FLD. (c) In the presence of read length column artifacts, the inferred FLD deviates from the true FLD, and exhibits a “phantom peak” at read length. (d) In the presence of read length missing-column artifacts, the inferred FLD deviates from the true FLD, and exhibits a “phantom peak” at read length.

### Local fragment length distribution varies around read length artifacts

In addition to binding events, we have observed local artifactual patterns that create dependence between fragment length and genomic position on both strands, preventing the reconstruction of fragment length distribution. These artifactual patterns fall into the following two categories:

- a pile of reads whose start positions constitute a read length width column pattern on the positive strand, followed by a read length width column pattern on the negative strand, a read length downstream. We refer to this artifact as a column artifact. We simulate it in Figure 3c, and show it in Figure 4a.
- a binding peak (or background read coverage) but with a read length width column pattern of missing reads on the positive strand followed by a read length width column pattern of missing reads on the negative strand, a read length downstream. We refer to this as a missing-column artifact. We simulate it in Figure 3d and show it in Figure 4b.

Both of these artifacts cause local disturbances in the fragment length distribution, however, the aggregate global fragment length distribution remains constant. This is easier to see in paired-end sequenced reads. The paired-end column artifact shown in Figure 4c contains fragments with length distributed according to the sample’s fragment length distribution. However, the fragments are organized such that longer fragments extend further from center of the artifact (denoted with an asterisk) than shorter fragments, implying *F* is dependent on *R^n^* and *R^n^*. The paired end missing-column artifact shown in Figure 4d is composed of fragments organized such that the range of possible fragment lengths is restricted based on genomic position. Specifically, on the positive strand, lengths of fragments (such as fragments A and B) must lie outside the range [*d^p^, d^p^* + *w*] where *d^p^* is the distance *d^p^* from the positive strand read to the center of the missing-column artifact (denoted with an asterisk). Likewise, on the negative strand, the lengths of fragments (such as fragments *C* and D) must lie outside the range [*d^n^, d^n^* + *w*] where *d^n^* is the distances *d^n^* from the negative strand read to the center of the missing-column artifact (denoted with an asterisk). Thus, column and missing-column artifacts are composed of fragments organized such that both strands are dependent on fragment length.

**Figure 4:**
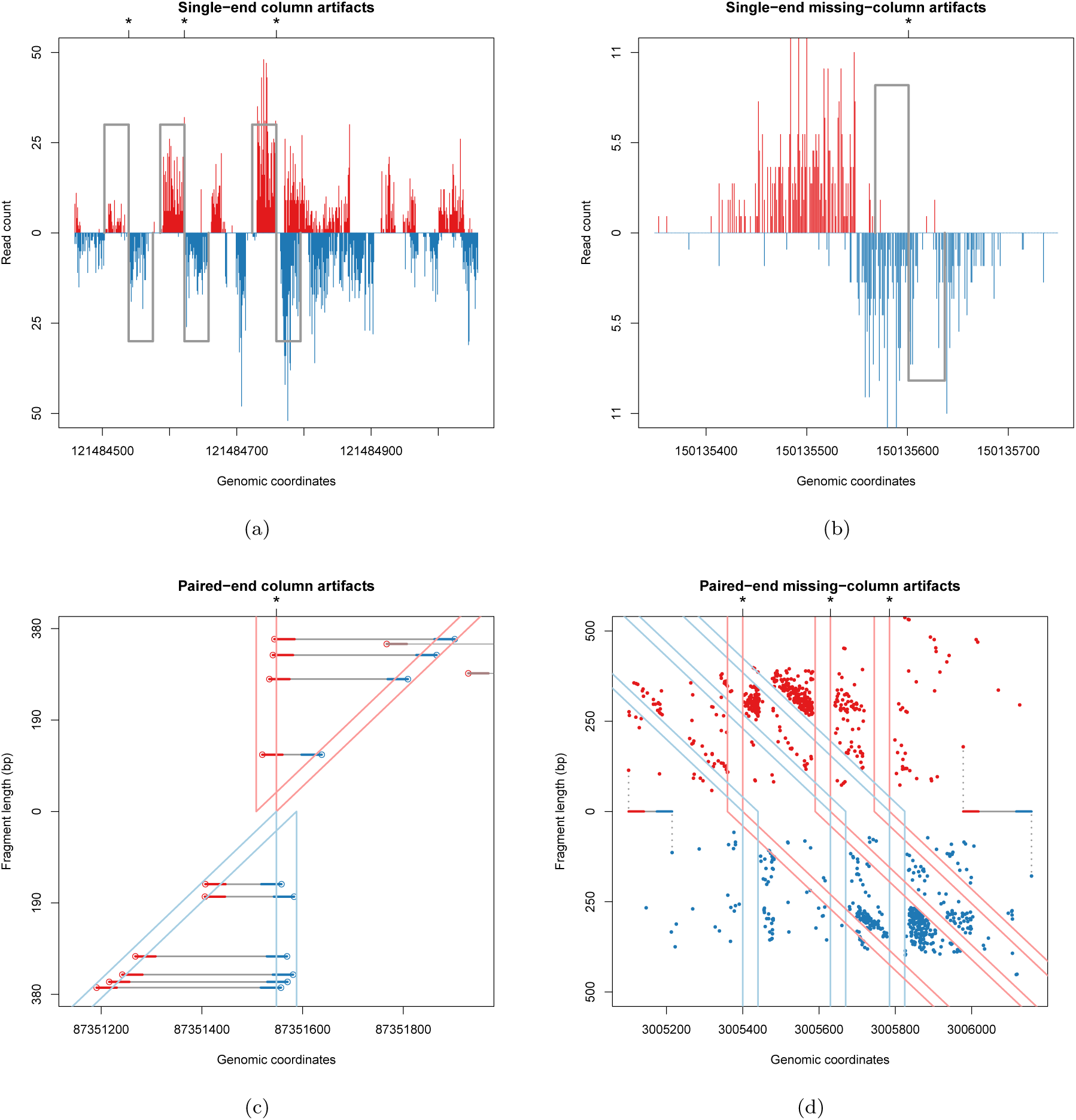
Column and missing-column artifacts and the nonrandom fragment length distribution in their neighborhoods. Examples of read length column (a) and missing-column (b) artifacts in a single-end human GM12878 cell anti-SRF ChIP sample on chromosome 1. Examples of a read length column (c) and missing-column artifact (d) in a paired-end MEF input control sample on chromosomes 1 and 18 respectively. Positive strand reads are shown in red while negative strand reads are shown in blue. The artifact center positions *x* where the sample genome differs from the reference are marked with asterisks. The paired-end scatterplots show each read’s position (x axis) and associated fragment length (y axis) to demonstrate the dependence relationship between genomic position and fragment length. Every positive strand read (red point) is accompanied by a negative strand read (blue point) originating from the same fragment. For clarity, in the paired-end column artifact plot (c), we have plotted each fragment and separated them to two groups such that in one group all fragments have positive strand reads within the positive strand column (light red column) and in the other group all fragments have negative strand reads within the negative strand column (light blue column). Explicitly, reads from fragments contributing positive strand columns are shown between the pink lines, while those from fragments contributing to the negative strand column are shown between cyan lines. The paired-end missing-column artifact (d) has a column of missing reads on the positive strand followed by a column of missing reads on the negative strand. The positive strand column of missing reads is linked to a diagonal of missing reads on the negative strand, representing the associated missing downstream fragment ends, and are together outlined in light red. Similarly, the negative strand column of missing reads is linked to a diagonal of missing reads on the positive strand, representing the associated missing upstream fragment ends, and are together outlined in light blue. We highlight two fragments to demonstrate the coupling between reads on the positive strand and reads on the negative strand.

We note that if we were using **Equations** 2 or 4 without invoking **Equations** 5 - 8 at these artifactual regions, the resulting fragment length distribution would have two modes, one associated with the “phantom peak” near the read length, and one associated with the predominant fragment length. ENCODE observed these two modes in the opposing strand cross-correlation, which is related to the fragment length distribution (**Equations** 1 and 3). However, when these artifactual regions are filtered out as is done in **Equations** 5–8, the “phantom peak” is greatly attenuated. Thus, these artifactual areas give rise to a low quality RSC [2, 49], the ENCODE measures of the “phantom peak” using cross-correlation.

### Incorrect mapping leads to read length artifacts

The read length used in any sequencing experiment is determined subsequent to the collection of fragments. Therefore, read length artifacts are associated with sequencing or post-sequencing procedures. Each read in a ChIP-seq experiment is sequenced from the sample’s genomic DNA, which can differ from the reference genome used in the alignment step. Comparative genome assembly algorithms used for sequence alignment work by comparing the nucleotide sequence for each read to the sequence of the reference genome, and assigning the read to the location that gave the best alignment score, usually based on fuzzy string matching. If the nucleotide sequence of the sample’s genome, from which the reads are sampled, disagrees with the reference genome at location *x*, the mapping algorithm can fail to assign the appropriate reads to that location. This could occur if the number of mismatched bases per read exceeds a predetermined cutoff or simply if the reads belonging to that location map better to another region with higher sequence similarity. When this occurs there will be a discontinuity in coverage across *w* nucleotides (where *w* is the read length) because there are exactly *w* possible read start positions where the read would overlap the mismatched base. On the positive strand the discontinuity is upstream of *x* and on the negative strand the discontinuity is downstream of *x*. This results in an missing-column artifactual coverage pattern of incorrectly mapped reads to the upstream sequence highlighted in yellow as seen in Figure 5. Further, reads that fail to map to the correct location can instead map to another location with higher sequence similarity as determined by the mapping algorithm. This would result in a column artifact as seen in the downstream sequence highlighted in yellow in Figure 5. These artifacts tend to occur in interspersed repetitive regions such as the sequences shown in yellow in Figure 5.

**Figure 5:**
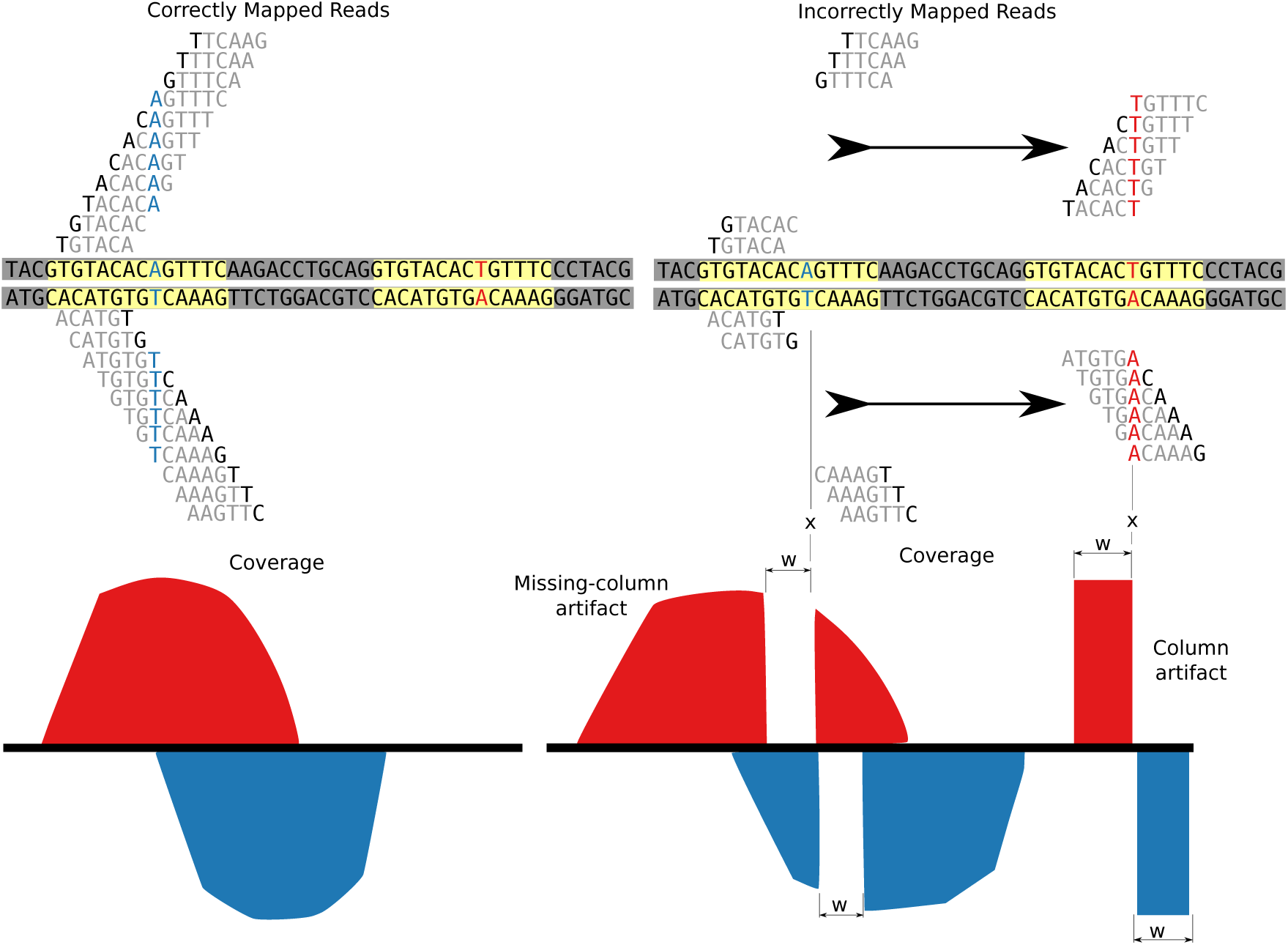
Read length artifacts likely stem from mapping problems. Reads that map to their correct locations do not give rise to artifacts (left). Differences between the reference and sequenced genomes can produce missing-column artifacts, and additionally the coverage can be relocated to a region of higher sequence similarity forming a column artifact (right).

For paired-end each relocated read in a column artifact, the associated read from the same fragment is also relocated as shown in Figure 4c. Likewise for each missing read in a missing-column artifact, the associated read from the same fragment is also missing as shown in Figure 4d.

### Derive the expected read coverage distribution around binding events

Once we infer the fragment length distribution, we use it to derive a filter matched to the read coverage pattern characteristic to regions of true binding events (step 3 of Figure 1). We denote putative ChIP target binding sites by *B_j_*, where *j* is an index representing the *j*-th putative binding event along the genome. Each fragment originating from binding event *j* covers the binding position, *B_j_*. The binding position *B_j_* is then related to the read position as follows:

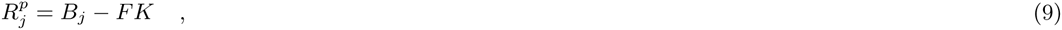
and

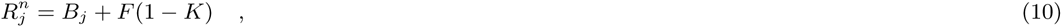
where 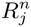 is the start position of a read on the positive strand resulting from event *j*, and 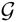 is the end position of a read on the negative strand resulting from event *j. K* is a random variable taking values between zero and one, describing the relative position of the binding site within a fragment. If *K* equals 0, then the binding position is at the most upstream end of the fragment, and if *K* equals 1, then the binding position is at the most downstream end of the fragment. If *K* is between 1 and 0, the binding position is at that location relative the fragment length. We model *K* as a beta distributed random variable Pr[*K*] ~ *B*(*α, β*). The beta distribution is convenient for this purpose because it is flexible for modeling random variables which take values between 0 and 1. Additionally, we set *α = β*, assuming that *K* is symmetrically distributed. We initialize *K* to a uniform distribution (*α* = 1) as a natural choice in the absence of prior knowledge (step 3 of Figure 1), and as detailed subsequently we reevaluate it by optimizing *α* (step 7 of Figure 1). Next, applying algebra of random variables, it can easily be seen that:

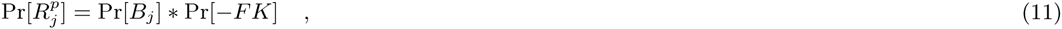
and

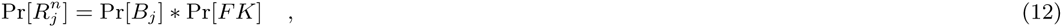
where * is the convolution operator, Pr[*K*] = Pr[1 − *K*] due to the symmetry mentioned above, and Pr[*FK*] is the product distribution as follows:

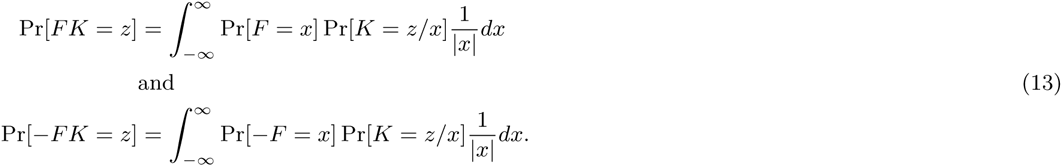

Pr[−*FK*] is the distribution of local read coverage on the positive strand with support upstream of a binding position (negative offset). Pr[*FK*] is the distribution of local read coverage on the negative strand with support downstream of a binding position (positive offset). We will use these local coverage patterns in both steps 4 and 6 of Figure 1 to locate and quantify candidate peaks respectively.

### Matched filtering for rapid and accurate localization of candidate peaks

Transcription factors bind to a small fraction of the genome, thus to improve efficiency we only test candidate peak positions (step 6 of Figure 1) that closely match our expected peak shape (Pr[−*FK*] and Pr[*FK*]), as identified by a matched filter [50] (step 4 of Figure 1).

In signal processing, a filter is a function which selects for the desired output signal vector, *s*, and suppresses the undesirable noise vector, *v*, of an observed input signal *x* = *s* + *v*. A matched filter [50] is a specialized filter whose time inverse is the impulse response function, *h*, where *h* is optimally parallel to the desired signal (*h || s*), and orthogonal to the noise (*h* ⊥ *v*). The matched filter has the favorable property that when convolved with an observed signal, it will maximize the output signal to noise ratio. The time inversed matched filter, *h*, is defined as follows:

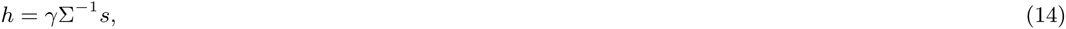
where *γ* is a normalization constant and Σ^−1^ is the inverse covariance matrix of the noise. For identifying peaks in ChIP-seq, the desired output signal *s* has shape Pr[−*FK*] for the positive strand and Pr[*FK*] for the negative strand.

Noise is rarely globally stationary (i.e. invariant probability distribution with respect to genomic position). However, when the noise distribution changes on a much larger length scale than the support (*2F_max_*) of the desired signal, it is fair to assume that the noise is independent and locally stationary, as is done in many other algorithms. Under this assumption, Σ, the covariance matrix of the noise at each position, is diagonal and locally proportional to the identity matrix. This property implies that in **Equation** 14 *h* ∝ *s*. Absorbing the normalization constants, the impulse response functions for the positive and negative strands can then be written as:

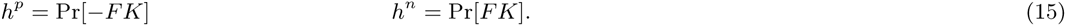

As a result the filtered signal *Y* at each position is given by the sum:

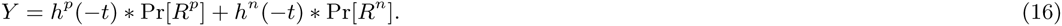

Usually *γ* is chosen to normalize the expected power of the noise after application of the filter to one. However, a given *γ* that normalizes the noise in low coverage areas to one, necessarily will give higher power in areas of higher coverage. Therefore, it is infeasible to specify a single *γ* when the noise is not globally stationary. As a result, the standard way of detecting the desired output signal by thresholding using a fixed signal to noise ratio is not applicable. Alternatively, we identify local maxima using a Gaussian derivative filter, a technique commonly used for detecting local maxima (edges) in images as follows:

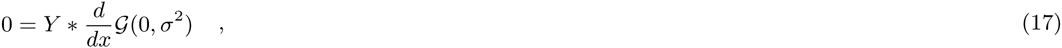
where 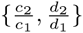 is the Gaussian distribution with variance *σ*^2^. Zero crossings (**Equation** 17) from positive to negative of this smoothed first derivative are the local maxima. A minimum read count requirement is applied to avoid spurious low coverage local maxima. Those local maxima passing the threshold are selected as candidate binding events, *b_j_*; *j* ∈ [1, *N*] (for *N* candidate events).

### Remove false positive binding events using cross-correlation

Genomic regions that have similarly high read coverage in both the ChIP sample and the negative control are false positive binding events. Most current peak calling algorithms rely on negative controls (usually total DNA input) to control the false positive rate. To the best of our knowledge, if negative controls are not available, false positive events would not be filtered out by current peak calling algorithms. We have discovered that most of the significant false positive events are in fact the read length artifacts described earlier and exemplified in Figure 6. We have already shown that the cross-correlations and deconvolved fragment length distributions of regions with read length artifacts exhibit the “phantom peak” at read length (Figure 3).

**Figure 6:**
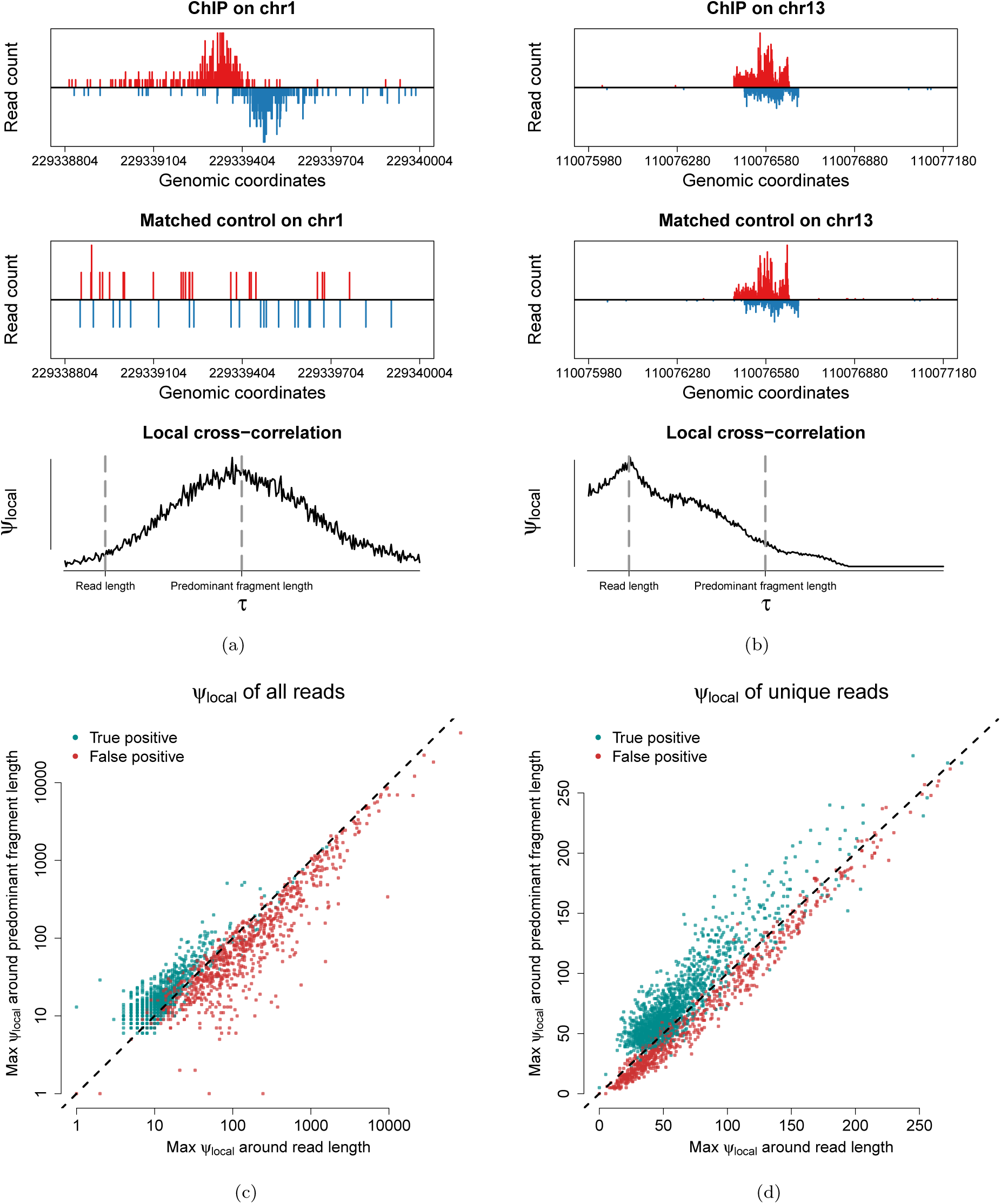
Local cross-correlation differentiates true binding events from read length artifacts (false positives). (a) Local cross-correlation of a binding event in anti-ATF2 K562 ChIP sample peaks near the average fragment length. (b) Local cross-correlation of a column artifact in anti-ATF2 K562 ChIP sample peaks near the read length. (c) A scatterplot of the maximum local log cross-correlation up to 10 bp beyond the read length versus the maximum local log cross-correlation in the range of 10 bp beyond the read length to 0.75F_*max*_ (d) A scatterplot of the maximum local binarized cross-correlation (using unique reads) up to 10 bp beyond the read length versus the maximum local binarized cross-correlation in the range of 10 bp beyond the read length to 0.75F_*max*_.

Ritornello identifies regions containing false positive events by detecting the “phantom peak” in their cross-correlations; true binding events contain no such “phantom peak” in thier cross-correlations. To this end, we employ a machine learning approach, building a classifier to distinguish between read length artifacts and true binding events. To extract features for the classifiers, we calculated cross-correlation locally from *b_j_ − F_max_* to *b_j_* + *F_max_* around each candidate peak. The features include: a) the maximum value of the cross-correlation function in the range between zero and read length, which is denoted by *c*_1_, and b) the maximum value of the cross-correlation function in the range between the read length and the maximum fragment length *F_max_*, which is denoted by *c*_2_. In the neighborhoods of binding events *c_2_* is expected to be higher than *c*_1_, whereas in neighborhoods of read-length artifacts we expect *c_1_* to be larger than *c*_2_. We added additional features to account for consecutive read length artifacts of varying amplitudes and large amplifications such as due to PCR. For this purpose, we binarized the coverage in the positive and negative strands by setting positions with read count greater than 0 to 1. We then performed a running mean smoothing on the binarized coverage, calculated cross-correlation and extracted the following features: a) the maximum value of the binarized smoothed cross-correlation function in the range between zero and read length, which is denoted by *d*_1_ and b) the maximum value of the binarized smoothed cross-correlation function in the range between the read length and the maximum fragment length *F_max_*, which is denoted by *d*_2_.

We build a classifier using logistic regression, with features: 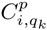. The instances used to build this classifiers include manually classified peaks obtained as follows: we first applied MACS2 (negative control free mode) to four transcription factor ChIP-seq datasets generated by ENCODE, subsequently selected the top 200 peaks for each of four samples, and finally manually labeled regions with typical binding shape as true positives (see Figure 6a) and regions with characteristic read length artifact as false positives (see Figure 6b). We trained this model using a five fold cross-validation and achieved high performance with AUROC of 0.993. This set of features is scale-free and thus our trained classifier is generalizable to any ChIP-seq sample and does not need to be retrained. Ritornello incorporates this trained classifier (step 5 of Figure 1) to flag artifactual locations as false positives without the need for a paired total DNA input or IgG control.

### Deconvolving single events from local coverage

The read coverage near an event is a mixture of reads generated by that event, as well as neighboring events and noise. In order to accurately quantify each event, it is essential to deconvolve its binding intensity (number of reads originating from each event) from this mixture. Fragments originating from different events in close proximity may overlap. Consequently, it is difficult to quantify the number of reads coming from each binding event. Further, fragments originating from non ChIP-ed DNA, off targeted sequencing due to antibody inefficiency, as well as other sources, contribute to background noise. We model the read coverage around each candidate peak using a generalized linear model to deconvolve its binding intensity.

The binding intensity, *β_j_*, of each candidate peak, *j*, is only dependent on positions where that peak has support, *i ∈* [*b_j_ − F_max_, b_j_* + *F_max_*]. To efficiently deconvolve the signal at *b_j_* we first discard peaks that do not overlap with *b_j_*. We retain only the subset *q_k_ ∈* {*q*_1_ *… q_T_*} of *T* peaks in close proximity to *j* (including *j*), such that the support of each peak in *q* overlaps with the support of *j*. The locally uniformly distributed noise associated with this neighborhood is indexed by *q*_0_. Here we assume that read counts follow a Poisson distribution, a common assumption made by other algorithms, such as MACS and GEM [3, 23]. We can then model the number of reads on the positive and negative strands, 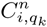 and 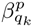, at position *i* due to event *q_k_* as follows:

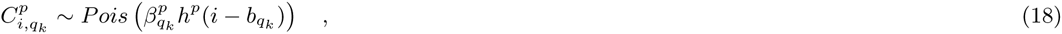
and

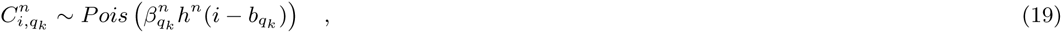
where the parameters 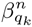 and 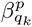 denote the binding intensities (expected read counts) of event *q_k_*. The impulse response functions *h^p^*(*i − b_qk_*) and *h^n^*(*i − b_qk_*) are the probabilities of observing a read at position *i* from event *q_k_*. We note that different 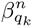 and 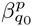 values are used to account for local differences in read coverage between positive and negative strands.

To model the noise we will once again invoke our assumption of locally stationary noise, as in the discussion before **Equation** 15. Here we assume that the locally stationary noise is a uniformly Poisson distribution. We model the read counts due to noise at position *i* for the positive and negative strands as follows:

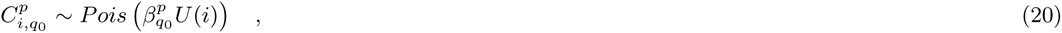
and

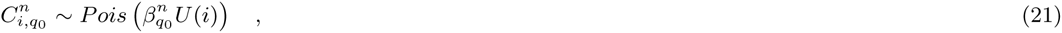
where *U* is a function that is locally uniform with support of 2*F_max_* around *b_j_*, and 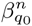 and 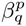 are the expected number of reads due to noise on the positive and negative strands respectively.

The read count at position *i* is then given by the sum of read counts from all sources *q_k_* as follows:

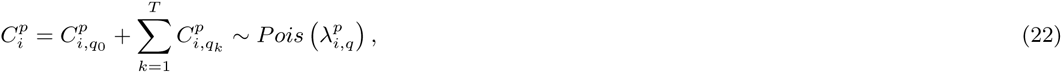
where

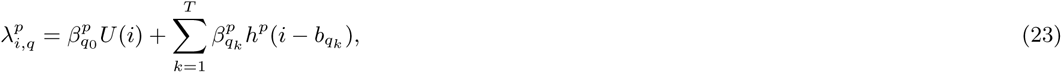
and

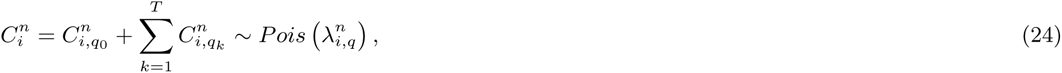
where

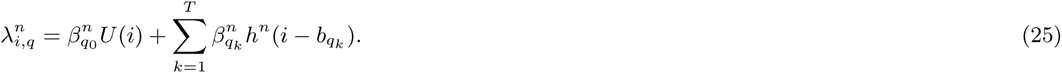

The relationships in **Equations** 22 and 24 use the following theorem: if *X*_1_ *… X_n_* are independent Poisson distributed random variables, *Pois*(*λ*_1_) *… Pois*(*λ_n_*), then their sum X_1_ + …+X*_n_* is Poisson distributed, *Pois*(*λ*_1_ *+* … + *λ_n_*).

In order to obtain the binding intensity for peak, *b_j_*, we maximize the likelihood for the models (**Equations** 22 and 24) of all nucleotides around *b_j_*. The likelihood of local binding intensities 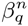 and 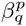 around the peak of interest, *j*, can be written as:

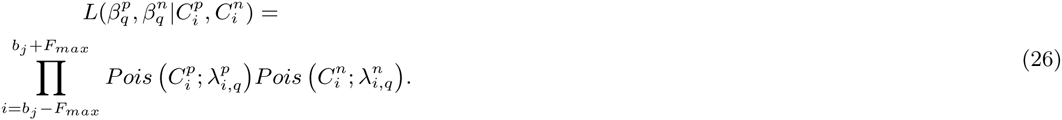

We then find the maximum likelihood estimates for parameters 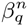 and 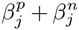. The sum, 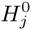, is reported as the binding intensity for the peak at *b_j_*.

We note that this is formally a Poissson generalized linear model with identity link function. Such a model has the advantage that it can resolve multiple peaks in close proximity, such as double binding or triple binding events. To our knowledge only BRACIL [46] and CSDeconv [47] are designed to deconvolve adjacent binding events, in particular double binding events. These algorithms are inefficient and therefore require as input a set of peaks from other peak callers. Ritornello implements a dogleg optimization (the Newton-Raphson method coupled with initial gradient descent), which is much faster than traditional Estimation Maximization or Markov Chain Monte Carlo methods [51], enabling the rapid deconvolution of all loci detected in previous steps.

### Testing candidate peaks for significance

In the previous section we quantified the intensity, *β_j_*, of each candidate peak. Here we determine the significance of each of these candidate binding events using a likelihood ratio test based on the likelihood we derived in **Equations** 26. The null model 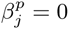 is obtained by setting both 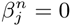 and 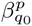. We note that we use the term null model at each position *b_j_* to refer to the model involving a zero binding intensity at *b_j_* but with potentially nonzero 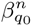 and 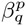 as well as 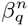 and 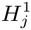 in neighboring candidate events. The alternative model 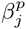 uses full parameterization including non vanishing 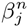 and 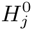 at the location of interest *b_j_*.

Since the null model 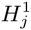 is nested within 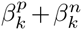, we can employ the likelihood ratio test statistic (D) in the form:

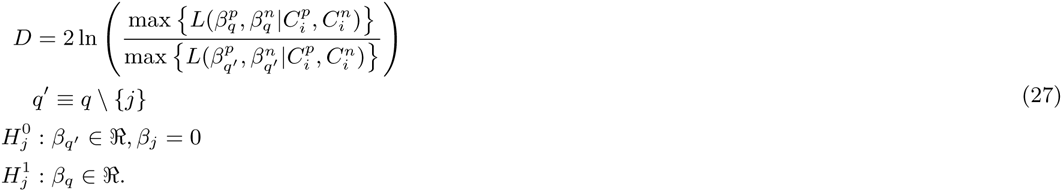

According to Wilke’s theorem [52] the likelihood ratio test statistic for this nested model is distributed according to a *χ^2^* distribution with two degrees. We then calculate the p-value for each peak based on this χ^2^ distribution. Finally, to account for false discovery in multiple hypothesis testing, we corrected these p-values using Benjamimi-Hochberg correction [53].

Up to this point, we have obtained an initial list of putative binding events. This was based on inferring an impulse response function given by the product distribution of the fragment length distribution and a uniformly distributed *K* (step 3 Figure 1). To further refine the impulse response function, we find the estimate of *α* that maximizes the combined likelihood of the 200 most significant putative events (step 7 of Figure 1). As shown in Figure 7, the Pr[−*FK*] and Pr[*FK*] derived from this procedure closely match the shape of highly abundant peaks. Finally, we repeat the peak identification, artifact testing, and likelihood ratio testing (steps 4–6 of Figure 1) using the updated *h^p^* and *h^n^* and report final list of significant peaks.

**Figure 7:**
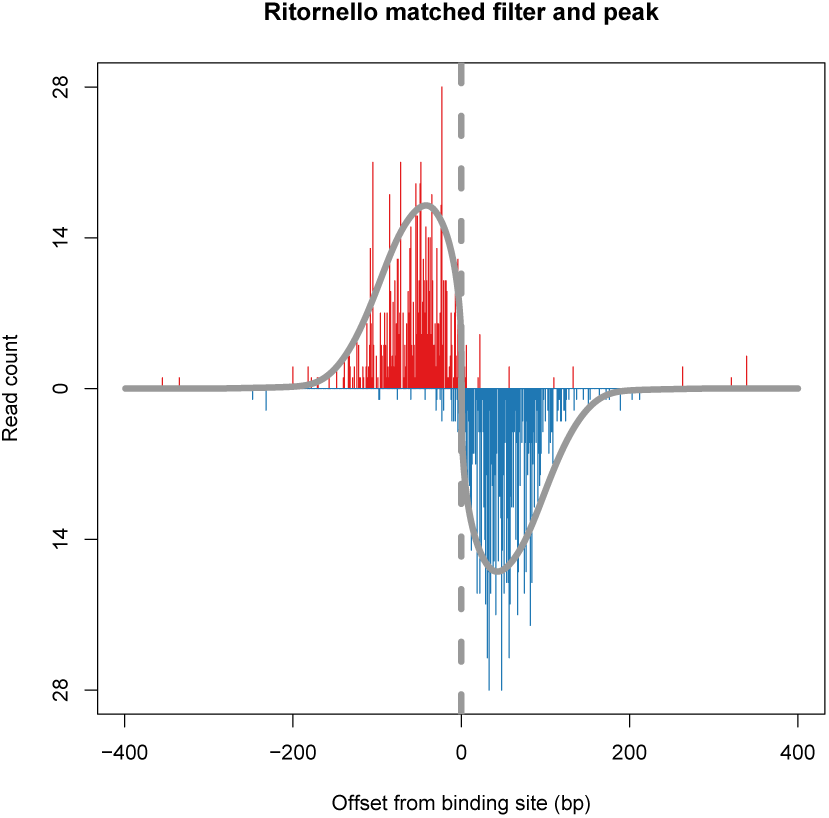
The parameterized filter closely matches the peak shape. Shown is an example peak in a human SRF sample. The parameterized filter for this sample is shown in grey. The peak location as determined by the filter is shown by a dashed line.

## Benchmarks

To assess Ritornello’s performance, we compared it against the MACS2 [3] and GEM [23] peak callers, which have both been recommended by the ENCODE consortium [54, 55]. We use 14 single-end transcription factor ChIP-seq experiments from the ENCODE project [55], each with two biological replicates (see Table 1 and Supplementary Table S1). Matched DNA input or IgG controls were also available (Supplementary Table S1). We run Ritornello, a control free algorithm, MACS2 (with the matched control), and GEM (with the matched control) on each of the 28 samples. We compare the performance of these algorithms in terms of:

**Table 1:** Transcription factor ChIP-seq experiments (two replicates each)

| TF | Cell type |
| --- | --- |
| NRF1 | K562 |
| SRF | GM12878 |
| REST | H1 |
| MAX | K562 |
| ATF2 | H1 |
| E2F4 | GM12878 |
| GATA1 | MEL |
| MYC | MEL |
| CTCF | Myotube |
| ELK1 | K562 |
| SRF | H1 |
| MAX | H1 |
| YY1 | H1 |
| REST | K562 |

- the degree of reproducibility between biological replicates
- the similarity between characteristic coverages in reproducible binding events predicted uniquely by Ritornello, MACS2, or GEM to the coverages of strong binding events predicted by multiple algorithms
- motif enrichment in top 1000 reproducible peaks

Additionally, we demonstrate that potential false positive events not predicted by MACS2 or GEM, but predicted if they run without a matched control, are generally not predicted by Ritornello. We also demonstrate that Ritornello predicts very few reproducible false positives in duplicated negative controls. This suggests that Ritornello has a low false positive rate.

### Comparing Ritornello and alternative methods based on reproducibility

We have evaluated the reproducibility between biological replicates of each algorithm using the Irreproducible Discovery Rate (IDR) approach [56]. The IDR method calculates the consistency of peak rankings in each replicate, and determines an optimal cutoff to report the reproducible peaks between replicates. It is recommended by ENCODE to analyze replicates of ChIP-seq experiments [55].

For each algorithm we applied the IDR approach and counted the number of reproducible peaks passing the recommended thresholds (IDR=0.05) for each the 14 experiments. We found that Ritornello had the largest number of reproducible peaks in 7 out of the 14 experiments, while MACS2 and GEM had a larger reproducibility in 5 and 2 out of the 14 experiments respectively (Figure 8). This indicates that Ritornello tends to capture similar or more reproducible binding events from biological replicates compared to MACS2 and GEM. This also demonstrates that the mathematical processing of the ChIP-seq signal can eliminate the need of using negative controls in TF ChIP-seq experiments. To evaluate the specificity of Ritornello, we applied the matched filter built in the ChIP channel to the matched control data and counted the number of reproducible false positives (values in parentheses in Figure 8). We found that Ritornello had few reproducible false positives in all five pairs of negative controls.

**Figure 8:**
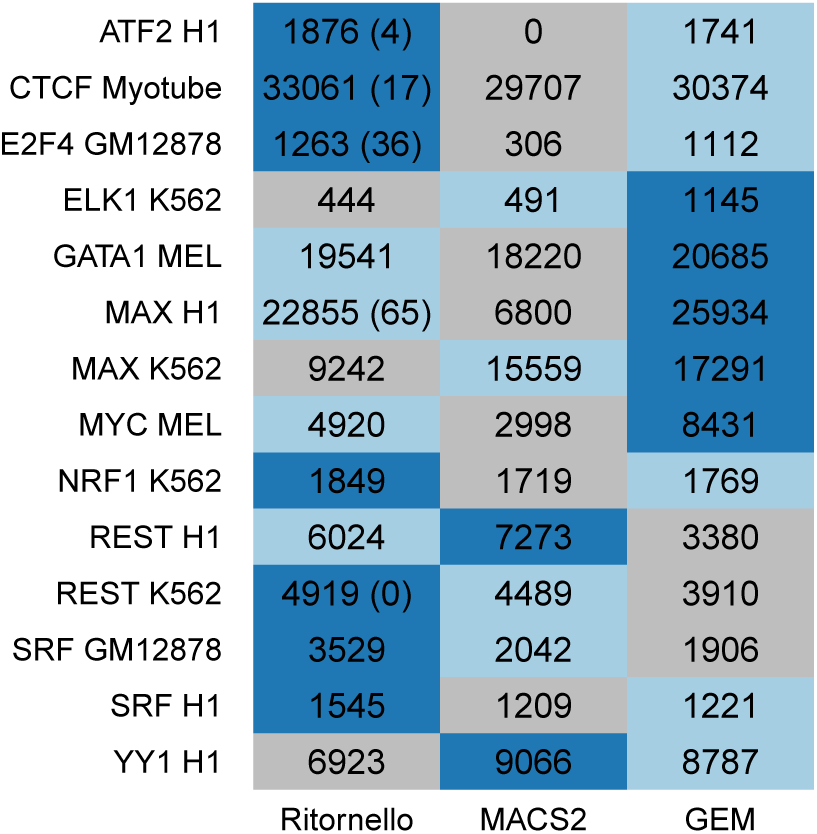
Number of reproducible peaks as determined by IDR between two biological replicates obtained by Ritornello, MACS2 and GEM. For each of the 14 experiments, we label in blue the algorithm that outputs the largest number of reproducible peaks, light blue the algorithm that outputs the second largest number reproducible peaks, and grey the algorithm that outputs the smallest number reproducible peaks. Five experiments out of the 14 presented have duplicated matched controls. Reproducibility of false binding in duplicated matched control (values in parentheses) demonstrates that Ritornello has a low false positive rate.

### Comparing Ritornello and alternative methods based on unique peak coverage patterns

To assess the shape and strength of reproducible peaks identified by the IDR approach, we compared the reproducible peaks reported by Ritornello to those reported by MACS2 and GEM respectively for each biological replicate. The majority of reproducible peaks reported by Ritornello, especially the most significant ones, overlap with those reported by MACS2 and GEM, indicating that Ritornello exhibits comparable performance to the other two algorithms even in the absences of negative controls (venn diagrams in Figures 9 and 10).

**Figure 9:**
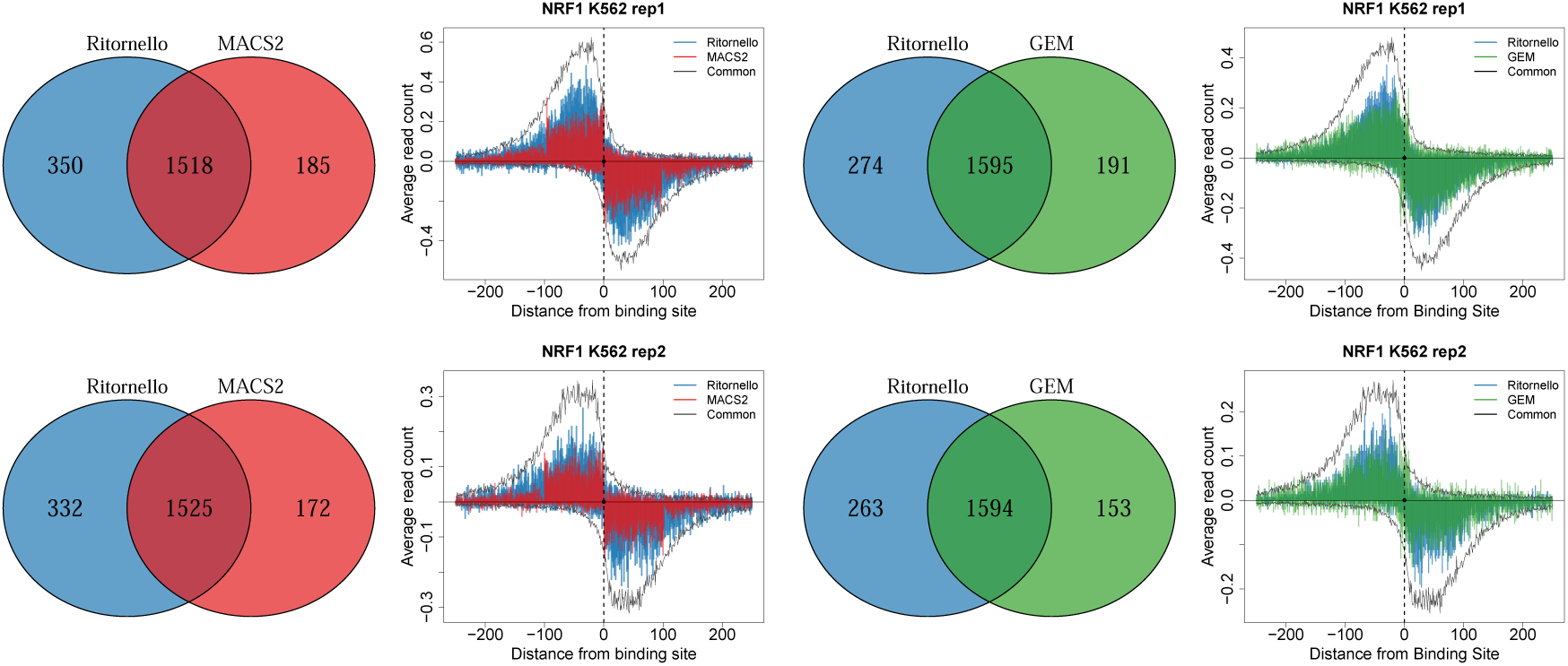
Pileup of read start positions for NRF1 reproducible peaks in K562 cells obtained by Ritornello only, by MACS2 only and by GEM only, where peaks within 100 bp of read length artifacts identified by Ritornello have been removed. The pileups of peaks common to Ritornello and MACS2 (left) or Ritornello and GEM (right) are shown in black. The pileups of read start positions for Ritornello best match the pileups of common peaks.

**Figure 10:**
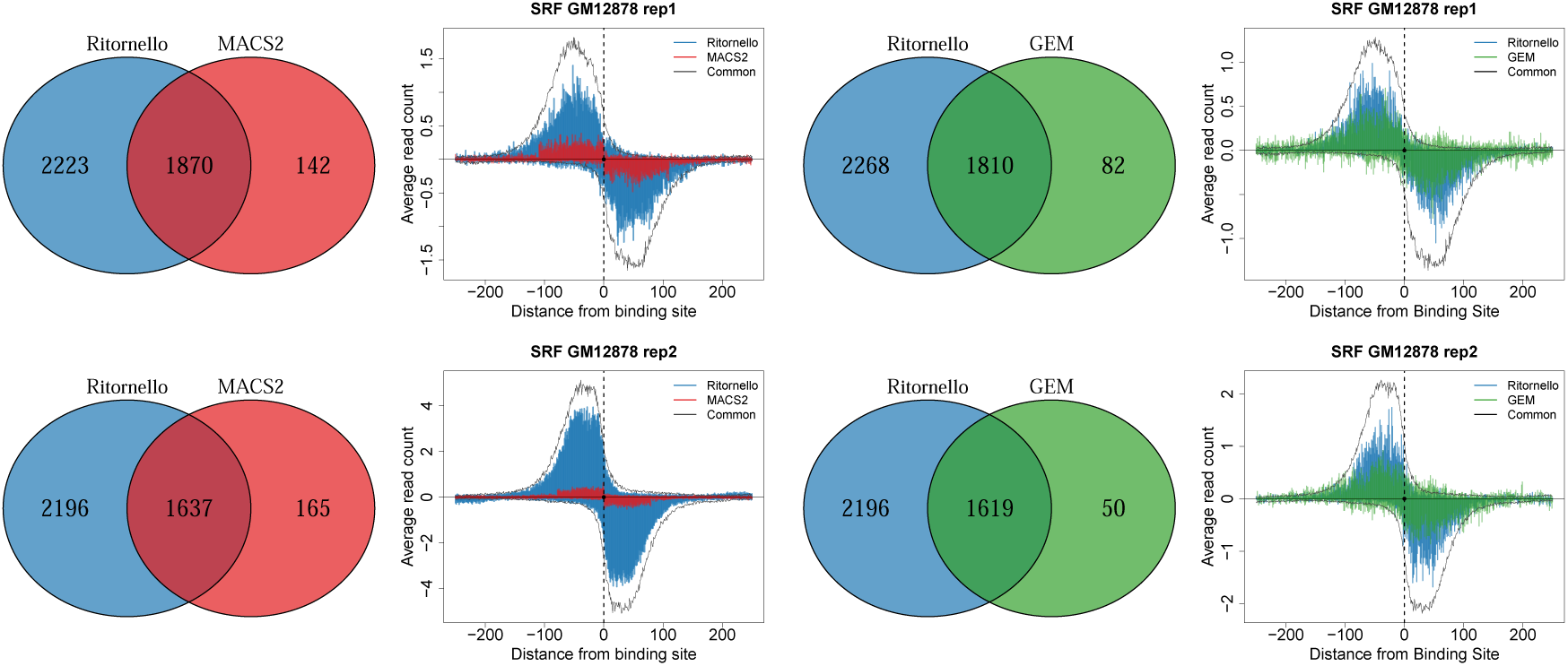
Pileup of read start positions for SRF reproducible peaks in GM12878 obtained by Ritornello only, by MACS2 only and by GEM only, where peaks within 100 bp of read length artifacts identified by Ritornello have been removed. The pileups of peaks common to Ritornello and MACS2 (left) or Ritornello and GEM (right) are shown in black. The pileups of read start positions for Ritornello best match the pileups of common peaks.

Further, we compared the reproducible peaks that are uniquely reported by Ritornello (but missed by MACS2) to those uniquely reported by MACS2. Similarly, we compared Ritornello-unique peaks to those uniquely reported by GEM. If we plot peaks reported by MACS2 or GEM, their pileup coverages are generally distorted by the aforementioned read length artifacts (Figures S1). In order to compare only those events in artifact free regions, prior to comparison we remove artifacts reported by Ritornello from both MACS2 and GEM reported peaks. For each comparison, we averaged the local distributions of read start positions (pileup) for the top 500 most significant unique peaks of each algorithm (pileup plots in Figures 9 and 10 and additionally Figures S2 - S13) and found that: a) peaks uniquely reported by Ritornello tend to have higher read counts around their binding sites than those uniquely reported by MACS2 or GEM, and b) the characteristic patterns (black curves whose shapes match the impulse response functions) associated with the highest intensity peaks reported by all algorithms tend to be similar to the patterns obtained by aggregating the pileups of peaks uniquely reported by Ritornello but in contrast are less similar to the aggregated pileups of peaks uniquely reported by MAC2 or GEM. We find this to be true even in those cases where GEM had similar read counts to Ritornello. These results imply that reproducible peaks uniquely reported by Ritornello are more likely to be true binding events than those uniquely reported by MACS2 or GEM.

### Comparing Ritornello and alternative methods based on motif occurrence rate

Availability of genuine validations of transcription factor binding events inferred by TF ChIP-seq peak callers is limited. Therefore, one of the measures used by practitioners for assessing the quality of peak callers is the fraction of predicted TF binding events that overlap with the characteristic binding motif of the relevant transcription factor. Employing the same 28 public ChIP-seq samples, we compare the motif enrichment for the top 1000 reproducible peaks reported by each algorithm as shown in (Figure 11). Predicted peaks whose binding centers are within 50 bp from the TF sequence motif are classified as events that are occupied by the motif. The peaks found by Ritornello had higher motif occurrence rate compared with MACS2 and GEM in 16 out of the 28 samples. MACS2 had higher motif occurrence rate compared with Ritornello and GEM in 6 out of the 28 samples, and GEM had higher motif occurrence rate compared with Ritornello and MACS2 in 5 out of the 28 samples. This suggests that Ritornello, which is a control free peak caller, is able to identify significant true binding events.

**Figure 11:**
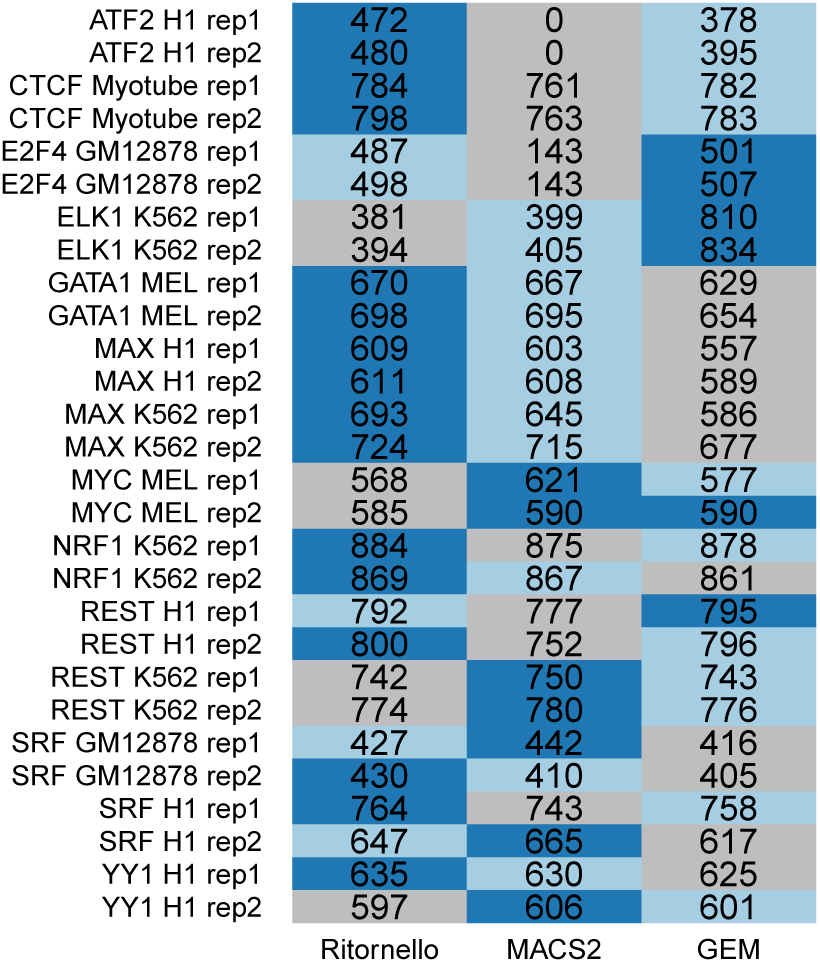
Occurrence of motifs within 50 bp of the top 1000 reproducible peaks obtained by Ritornello, MACS2 and GEM. For each of the 28 experiments (replicates are denoted by repl and rep2), we label in dark blue the algorithm that outputs the largest number of motif containing peaks, light blue the algorithm that outputs the second largest number of motif containing peaks, and grey the algorithm that outputs the smallest number of motif containing peaks.

### Ritornello identifies true binding events in low quality samples

ENCODE recommends discarding low quality samples as determined by the NSC and RSC scores. Ritornello discards read length artifacts that give rise to low RSC and uses a matched filtering approach that maximizes the signal to noise ratio measured by the NSC, we therefore conjectured that Ritornello may be able to rescue these low quality samples. We compared the performance of Ritornello with that of MACS2 and GEM on samples with suboptimal quality based on the NSC and RSC scores. The ENCODE Consortium has suggested repeating experiments with NSC values less than 1.05 and RSC values less than 0.8. Using these criteria we identified that out of the 28 samples we investigated, four samples have suboptimal quality. These four experiments include: ATF2 H1 replicate 1 (NSC=1.04,RSC=0.62), ATF2 H1 replicate 2 (NSC=1.04,RSC=0.74), ELK1 K562 replicate 1 (NSC=1.03,RSC=0.64), and ELK1 K562 replicate 2 (NSC=1.05,RSC=0.73). We observed that in these four samples, the pileups of reproducible peaks predicted by Ritornello have a characteristic bimodal shape of transcription factor binding and have much stronger read coverage than their matched input controls (Figure 12a). This demonstrates that there are numerous significant binding events that can be captured in low quality ChIP-seq samples. Additionally, the pileups of reproducible peaks reported by MACS2 or GEM have either uniform read coverage or a narrow bimodal shape (similar to column artifacts) as in Figure 12b. This illustrates that Ritornello reliably rescues peaks from low quality samples.

**Figure 12:**
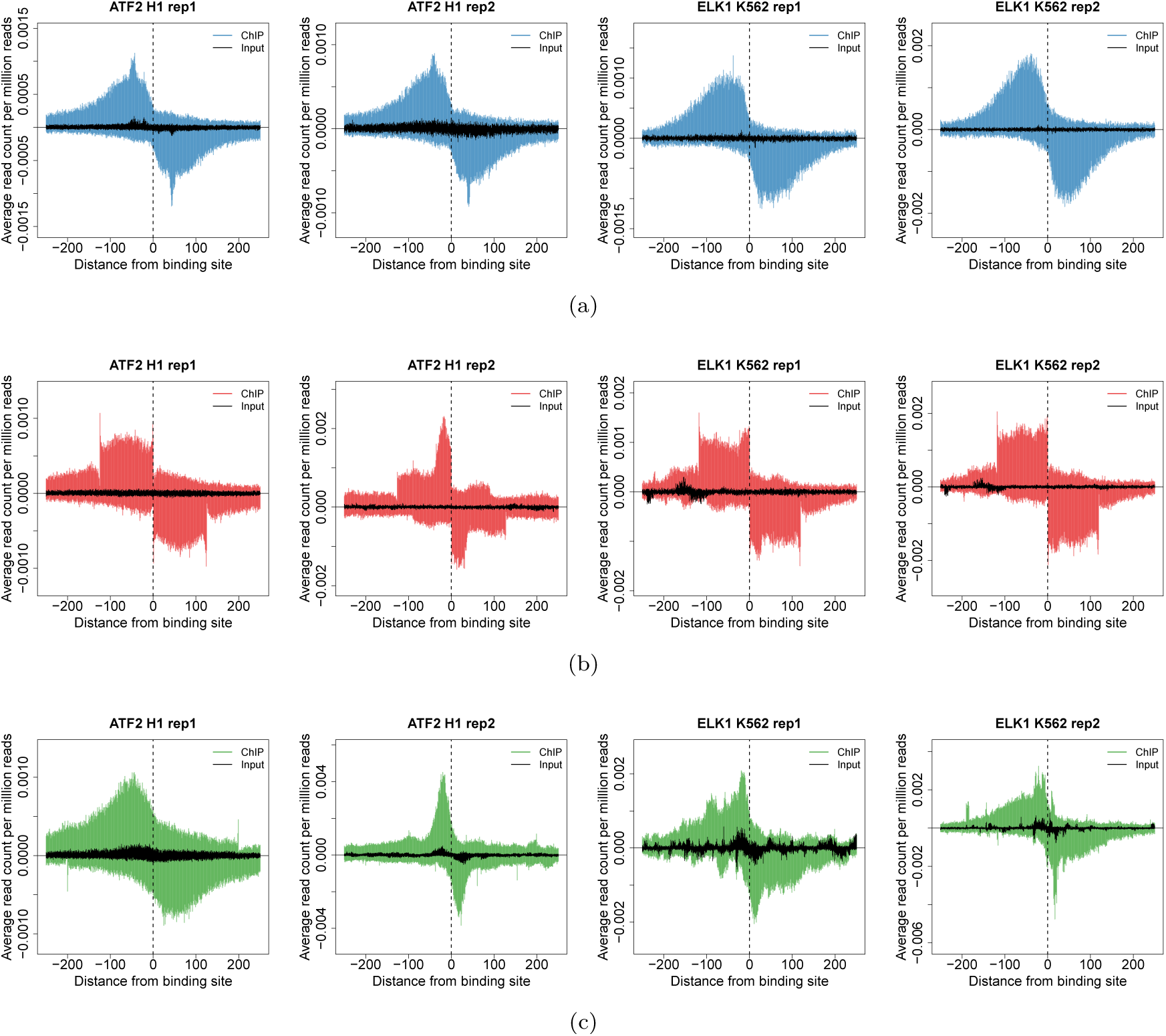
Pileup of read start positions for reproducible peaks identified by Ritornello, MACS2 and GEM in samples of low quality and their matched controls. (a) The pileups of reproducible peaks called by Ritornello in ChIP samples are shown in blue, and the pileups in the matched controls are shown in black. The pileups of peaks detected by Ritornello show smooth bimodal shapes and have much stronger read coverage in the ChIP as compared to their matched controls. (b) Pileups of read start positions from peaks detected by MACS2 are shown in red. The pileups of reproducible peaks called by MACS2 have a narrow bimodal shape (column artifact) or uniform coverage shape. (c) Pileups of read start positions from peaks detected by GEM are shown in orange. The pileups of reproducible peaks by GEM also have a narrow bimodal shape or irregular spike like in the negative controls.

### Ritornello obviates the need for a matched input control

Although matched input controls are optional for some methods, they are used by ChIP-seq peak calling algorithms to avoid calling many spurious false positives. Both MACS2 and GEM have options to run with or without the matched input control. Notably, Ritornello, which is a matched control free approach, avoids calling these false positives. To show this, we curated lists of potential false positives from each of the 28 samples. These lists were generated by considering the strongest peaks predicted by matched control free MACS2 (or GEM) but not predicted by MACS2 or GEM. To further enrich for false positives, the lists were subsequently filtered to exclude motif containing peaks. In most samples, Ritornello calls a small fraction of the false positive peaks (median=0.04) as seen in Table 2. This demonstrates that while standard methods such as MACS2 and GEM require matched control data in order to exclude false positives, Ritornello can discard most of these false positives without matched control data.

**Table 2:** Ritornello outputs a small fraction of peaks discarded by the use of a matched control. The proportion of motif-free reproducible peaks predicted by MACS2 (or GEM) without a matched control that are predicted by Ritornello but discarded by MACS2 (or GEM) when using matched control. This proportion is explicitly defined as 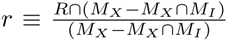 where *R* are the peaks reported by Ritornello, *M_X_* are the peaks reported by MACS2 (or GEM) without a matched control that do not overlap a motif and *M_I_* are the peaks reported by MACS2 (or GEM) with a matched control.

| | $r_{MACS2}$ | $r_{GEM}$ |
| --- | --- | --- |
| NRF1 K562 rep1 | 0.00 | 0.00 |
| NRF1 K562 rep2 | 0.02 | 0.03 |
| SRF GM12878 rep1 | 0.15 | 0.12 |
| SRF GM12878 rep2 | 0.03 | 0.02 |
| REST H1 rep1 | 0.03 | 0.02 |
| REST H1 rep2 | 0.03 | 0.02 |
| MAX K562 rep1 | 0.03 | 0.02 |
| MAX K562 rep2 | 0.04 | 0.09 |
| ATF2 H1 rep1 | 0.02 | 0.01 |
| ATF2 H1 rep2 | 0.33 | 0.20 |
| E2F4 GM12878 rep1 | 0.00 | 0.01 |
| E2F4 GM12878 rep2 | 0.19 | 0.09 |
| GATA1 MEL rep1 | 0.07 | 0.05 |
| GATA1 MEL rep2 | 0.08 | 0.07 |
| MYC MEL rep1 | 0.13 | 0.04 |
| MYC MEL rep2 | 0.15 | 0.05 |
| CTCF Myotube rep1 | 0.28 | 0.16 |
| CTCF Myotube rep2 | 0.06 | 0.05 |
| ELK1 K562 rep1 | 0.00 | 0.00 |
| ELK1 K562 rep2 | 0.00 | 0.00 |
| SRF H1 rep1 | 0.20 | 0.01 |
| SRF H1 rep2 | 0.02 | 0.00 |
| MAX H1 rep1 | 0.11 | 0.09 |
| MAX H1 rep2 | 0.27 | 0.18 |
| YY1 H1 rep1 | 0.00 | 0.02 |
| YY1 H1 rep2 | 0.02 | 0.02 |
| REST K562 rep1 | 0.01 | 0.04 |
| REST K562 rep2 | 0.11 | 0.09 |

We observed that in three samples (ATF2 replicate 2, CTCF replicate 1, and MAX hESC replicate 2 in Table 2), Ritornello had a non-negligible overlap with the potential false positive lists. To inspect whether the potential false positives that Ritornello reported are actual false positives, we created a pileup for these peaks. We observed in Figure 13, that these pileups closely resemble the most significant true binding event pileups, and are likely not false positives. This suggests that using MACS2 or GEM with a matched negative control may result in false negatives. Together, these results indicate that Ritornello both reduces the false positives in scenarios where standard methods are used without matched control data and detects putative events that standard methods using matched control data would fail to predict.

**Figure 13:**
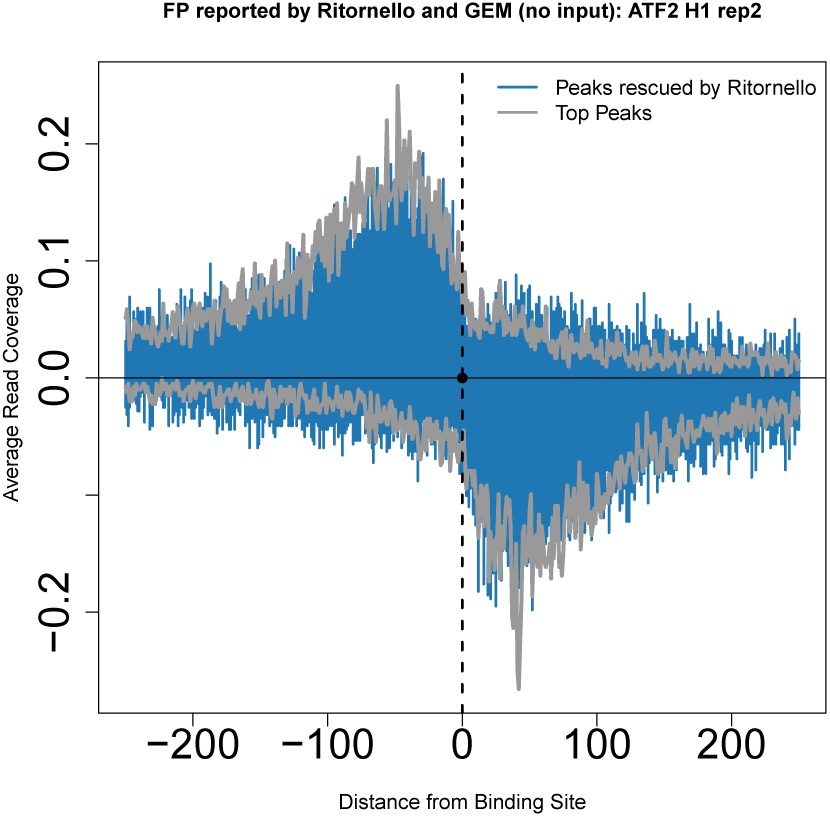
Ritornello rescues false negative events. Here false negatives are defined as reproducible peaks picked up by MACS2 or GEM without input, but discard by MACS2 and GEM with input. Ritornello also identifies these events in both biological replicates, which are likely to represent actual binding events because their pileups are similar to those of true binding events.

## Discussion

In this work, we demonstrated that we could infer the entire fragment length distribution, rather than only the mean fragment length, using a deconvolution approach from single-end TF ChIP-seq experiments. We derived an experiment specific probabilistic model to mathematically describe the well-known bimodal shape of TF binding. Using this bimodal shape, we applied the matched filter technique from signal-processing to identify potential TF binding sites and used a Poisson GLM to deconvolve the binding intensities and test the significance of each putative binding event. Our model efficiently deconvolves the effect of neighboring peaks as well as noise to resolve multiple adjacent binding events. We compared Ritornello (a control-free approach) with two popular algorithms recommended by ENCODE, MACS2 and GEM, which require matched controls to reduce false positives. We found that Ritornello outperforms these other methods in terms of reproducibility between biological replicates, motif enrichment of most significant reproducible peaks and the coverage pattern of unique reproducible peaks.

We also identified artifactual binding regions where the local cross-correlation peaks at read length instead of around fragment length. We elucidated that these artifactual regions contribute to the phantom peaks, associated with poor experimental quality. Current peak calling algorithms, such as MACS2 and GEM, rely on matched control samples to remove a substantial fraction of these artifacts. We provide an extensive description of this specific category of artifacts and their origin, and offer an automated approach to filter out artifacts without requiring matched controls. Taken together, Ritornello offers an alternative that obviates the need for a match control, demonstrating that one can safely reduce the total experimental cost of TF ChIP-seq experiments, while providing superior analytic results.

ENCODE provides a blacklist of genomic regions which contains artifactually high read coverage in different ChIP-seq experiments [57]. This manually curated blacklist largely overlaps with repetitive regions in the genome [57]. The blacklist has several drawbacks: 1) it does not cover all artifactual regions, 2) it is not generalizable to different cell types and 3) it is only available for human and mouse. We note that a few peak calling tools, such as PePr [58], optionally remove artifacts in regions where the local read coverage in the ChIP is similar to that in the matched control. However, these methods still require a matched control. Ritornello is capable of removing artifacts independently without requiring either prior knowledge of a blacklist or matched negative controls.

Many variables influence the quality of ChIP-seq experiments and our ability to infer true binding events from the data. These include factors such as antibody efficiency, DNA fragmentation, PCR amplification, sequencing depth, read mapping quality etc. Each of these factors may vary from one sample to another. Ritornello is designed to implicitly take into consideration these experiment specific parameters from raw data and is applicable to a wide variety of protocols. Additionally, it does not require any tuning of parameters. One limitation of Ritornello is that it is designed to detect point-source peaks such as TF binding events. For broad-source peaks, such as epigenetic modifications, we recommend other peak callers.

We demonstrated that in TF ChIP-seq experiments that would otherwise be discarded due to low quality, Ritornello reliably recovers true binding events. Often repeat experiments with the same reagents are of poor quality according to ENCODE’s metric and thus algorithms capable of handling such data are required. If, however, the repeated experiments were to improve results, the previous poor quality samples may still be of value to strengthen the findings of the higher quality samples.

### Software availability

Ritornello was programmed in C++ using the FFTW [59] library for fast computation of the Fourier transform and the Samtools [60] library for interfacing with the sequence alignment/map format which has become the standard in high throughput sequencing. Ritornello is freely available for download at https://sourceforge.net/projects/ritornello/files/. Further analysis and graphics were made using the R statistical language [61].

## Acknowledgements

The authors thank Roy Lederman, Vladimir Rokhlin, Ronen Talmon, and Ronald Coifman for insightful discussions.

---

**Table S1:** ChIP-seq samples

| TF | Cell type | ChIP | ChIP<br>Depth | Control | Control<br>Depth | Input/IgG |
| --- | --- | --- | --- | --- | --- | --- |
| NRF1 | K562 | ENCFF000YVE | 14422328 | ENCFF000YQR | 20216505 | IgG |
| NRF1 | K562 | ENCFF000YVG | 13140831 | ENCFF000YQR | 20216505 | IgG |
| SRF | GM12878 | ENCFF000OFD | 22677401 | ENCFF000ODH | 27578721 | Input |
| SRF | GM12878 | ENCFF000OFG | 31168812 | ENCFF000ODH | 27578721 | Input |
| REST | H1 | ENCFF000OQE | 15153099 | ENCFF000OSE | 66830473 | Input |
| REST | H1 | ENCFF000OQH | 20842565 | ENCFF000OSE | 66830473 | Input |
| MAX | K562 | ENCFF000YTD | 8120174 | ENCFF000YQR | 20216505 | IgG |
| MAX | K562 | ENCFF000YTJ | 20023038 | ENCFF000YQR | 20216505 | IgG |
| ATF2 | H1 | ENCFF000OME | 38468114 | ENCFF000OSH | 55951481 | Input |
| ATF2 | H1 | ENCFF000OMH | 42553524 | ENCFF000OSF | 27548706 | Input |
| E2F4 | GM12878 | ENCFF000VUG | 20373471 | ENCFF000VWA | 6211790 | IgG |
| E2F4 | GM12878 | ENCFF000VUH | 23847999 | ENCFF000VWB | 5526289 | IgG |
| GATA1 | MEL | ENCFF001NSH | 13589299 | ENCFF001NUG | 29225622 | IgG |
| GATA1 | MEL | ENCFF001NSI | 14533141 | ENCFF001NUG | 29225622 | IgG |
| MYC | MEL | ENCFF001NPX | 13253064 | ENCFF001NUD | 20400132 | IgG |
| MYC | MEL | ENCFF001NPY | 12649437 | ENCFF001NUD | 20400132 | IgG |
| CTCF | Myotube | ENCFF000BNJ | 13786817 | ENCFF000BNE | 42199165 | Input |
| CTCF | Myotube | ENCFF000BNL | 18693675 | ENCFF000BNF | 31202572 | Input |
| ELK1 | K562 | ENCFF000YMM | 20476073 | ENCFF000YQR | 20216505 | IgG |
| ELK1 | K562 | ENCFF000YMP | 14999686 | ENCFF000YQR | 20216505 | IgG |
| SRF | H1 | ENCFF000OUH | 25965316 | ENCFF000OSE | 66830473 | Input |
| SRF | H1 | ENCFF000OUM | 21984384 | ENCFF000OSE | 66830473 | Input |
| MAX | H1 | ENCFF000OPO | 28332084 | ENCFF000OSH | 55951481 | Input |
| MAX | H1 | ENCFF000OPR | 31783032 | ENCFF000OSF | 27548706 | Input |
| YY1 | H1 | ENCFF000OWG | 30394170 | ENCFF000OSE | 66830473 | Input |
| YY1 | H1 | ENCFF000OWI | 15877505 | ENCFF000OSE | 66830473 | Input |
| REST | K562 | ENCFF000QCL | 27965664 | ENCFF000QER | 13847077 | Input |
| REST | K562 | ENCFF000QCM | 26398569 | ENCFF000QES | 20248824 | Input |

**Figure S1:**
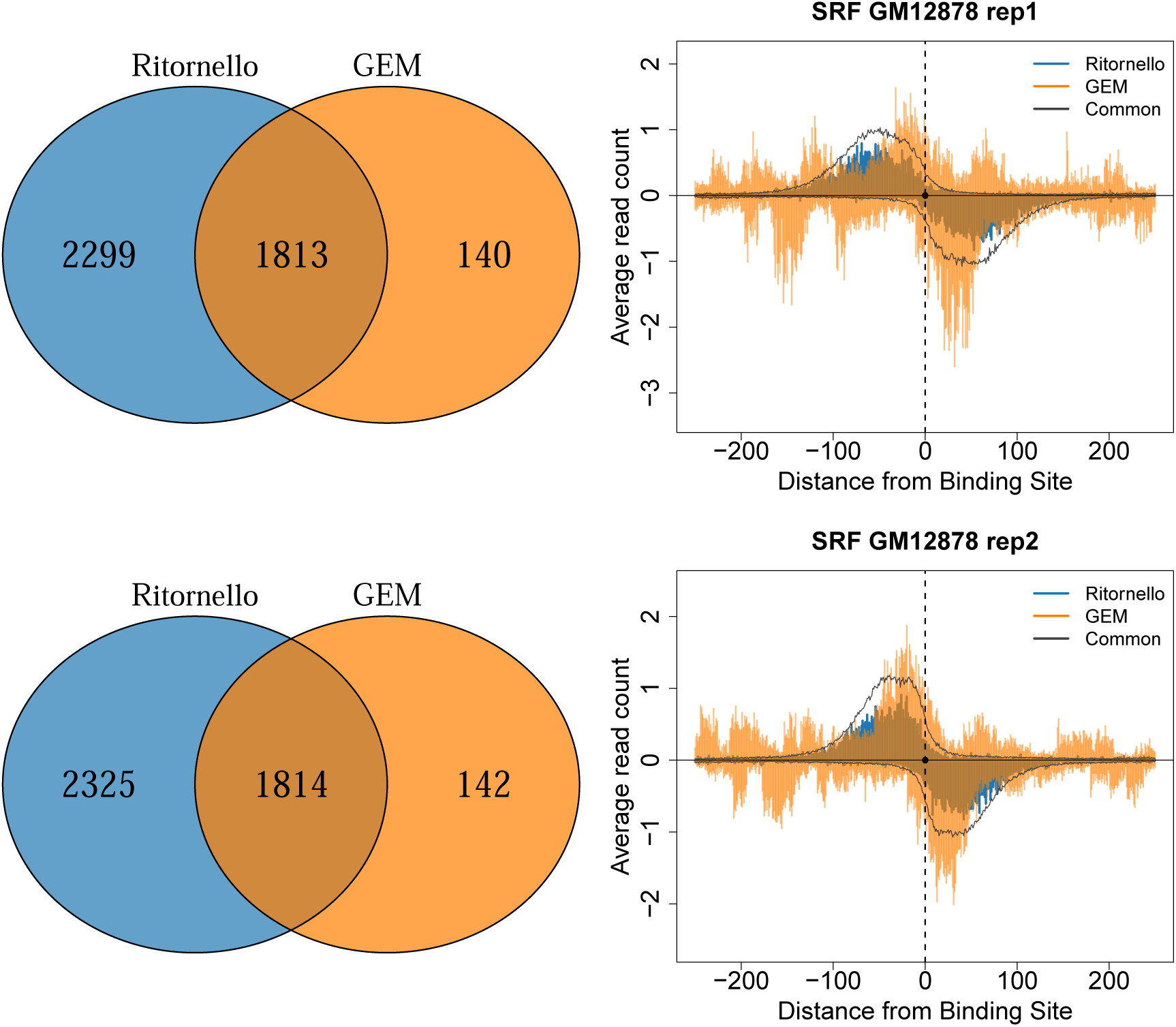
Pileup of read start positions for SRF reproducible peaks in GM12878 obtained by Ritornello only and by GEM only. Here all peaks called by GEM have been included (artifacts detected by Ritornello were not removed in contrast to Figure 10 where these artifacts have been removed). The pileups of peaks common to Ritornello and MACS2 (left) or Ritornello and GEM (right) are shown in black. Note that the number of peaks reported by Ritornello are different than in Figure 10 because there are a few peaks in close proximity to artifacts, and so when we remove peaks within 100 bp of artifacts we will remove these as well.

**Figure S2:**
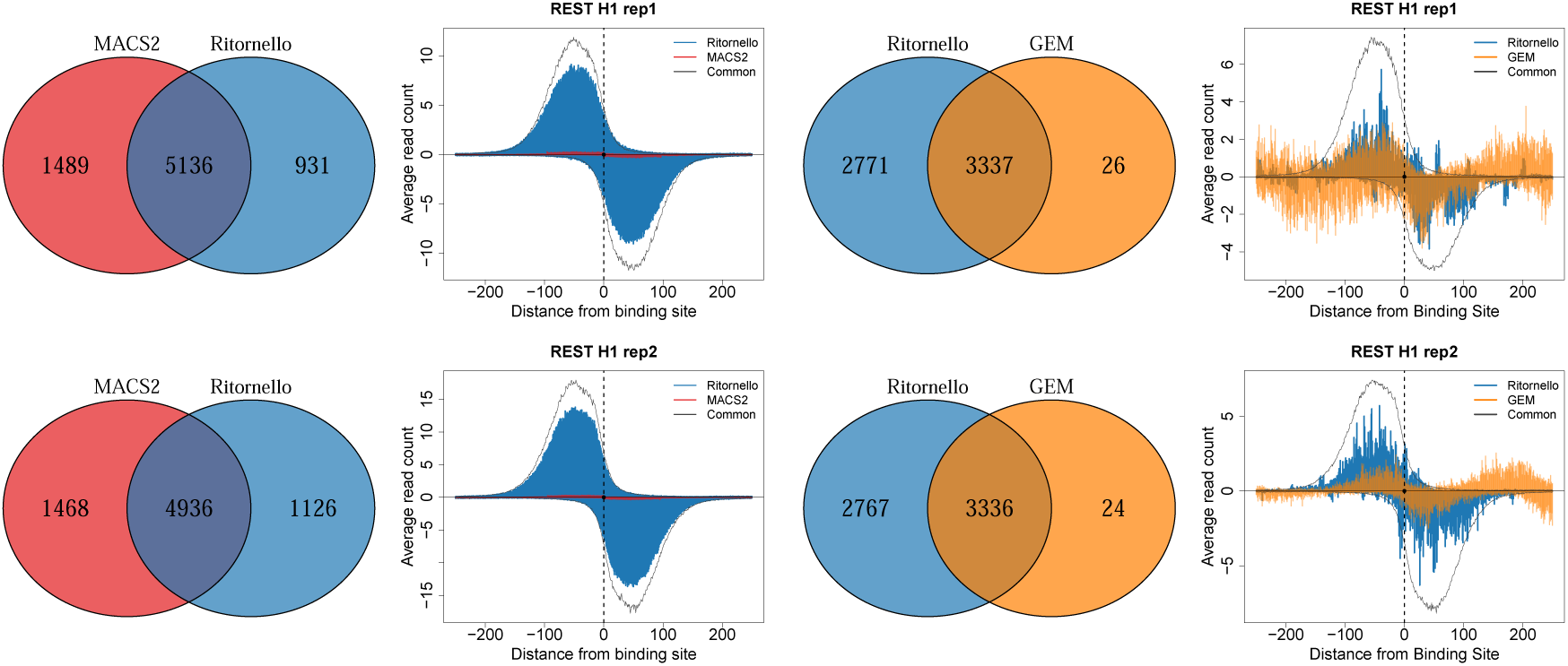
Pileup of read start positions for REST reproducible peaks in H1 obtained by Ritornello only, by MACS2 only and by GEM only, where peaks within 100 bp of read length artifacts identified by Ritornello have been removed. The pileups of peaks common to Ritornello and MACS2 (left) or Ritornello and GEM (right) are shown in black.

**Figure S3:**
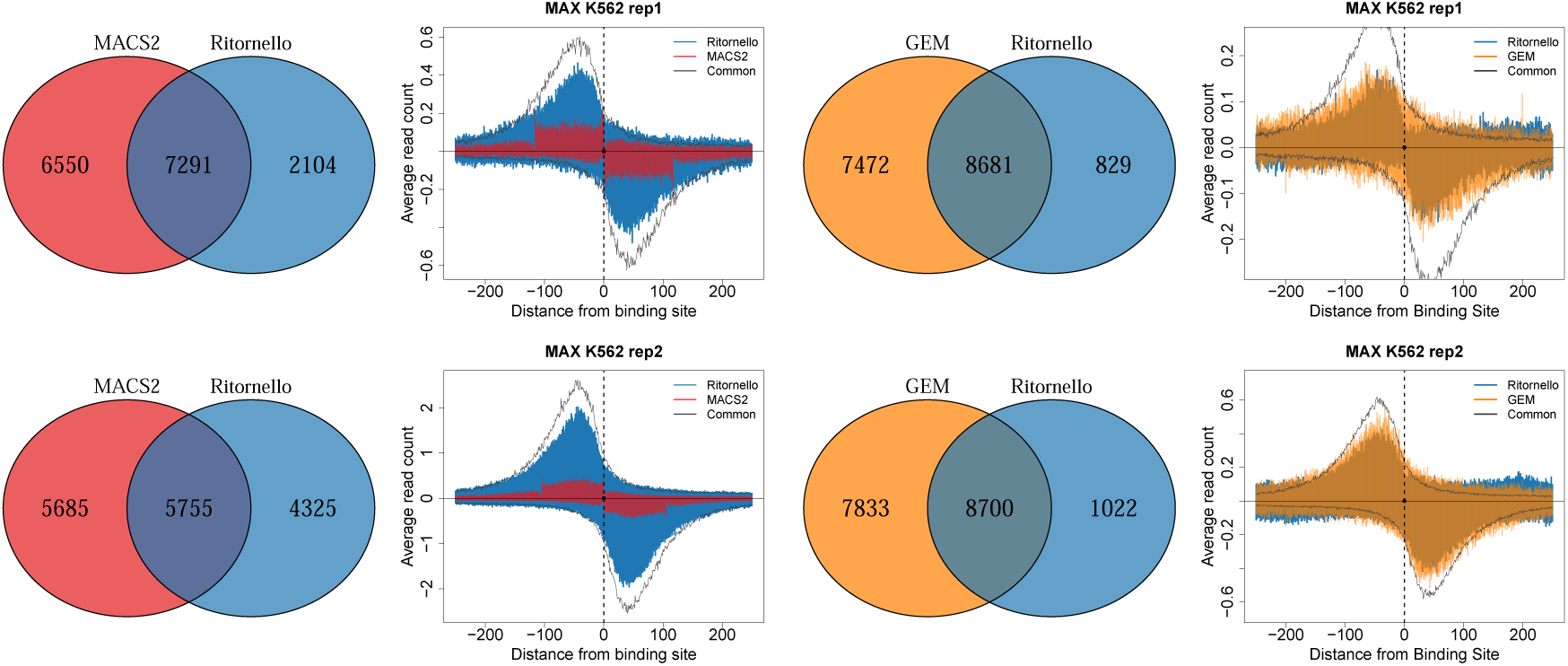
Pileup of read start positions for MAX reproducible peaks in K562 obtained by Ritornello only, by MACS2 only and by GEM only, where peaks within 100 bp of read length artifacts identified by Ritornello have been removed. The pileups of peaks common to Ritornello and MACS2 (left) or Ritornello and GEM (right) are shown in black.

**Figure S4:**
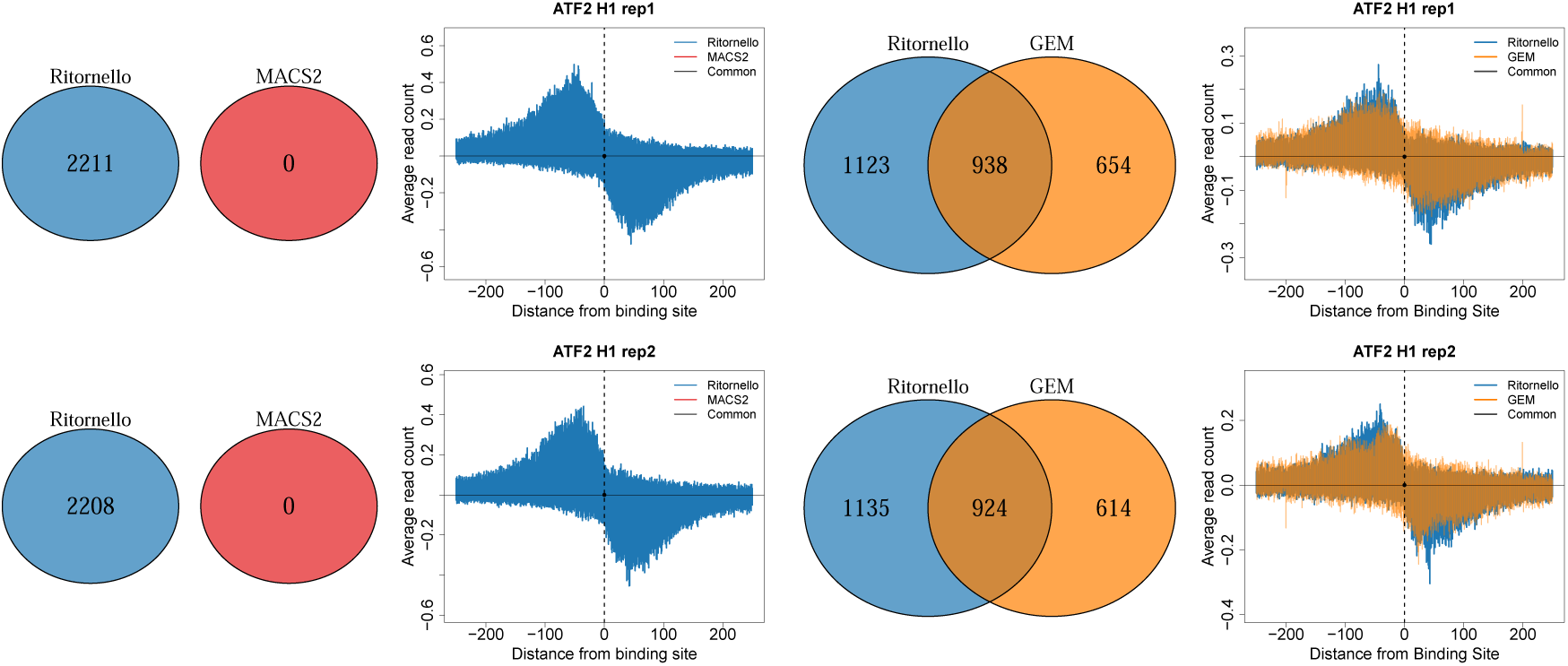
Pileup of read start positions for ATF2 reproducible peaks in H1 obtained by Ritornello only, by MACS2 only and by GEM only, where peaks within 100 bp of read length artifacts identified by Ritornello have been removed. The pileups of peaks common to Ritornello and MACS2 (left) or Ritornello and GEM (right) are shown in black. 1026 303 2 Ritornello MACS2 −200

**Figure S5:**
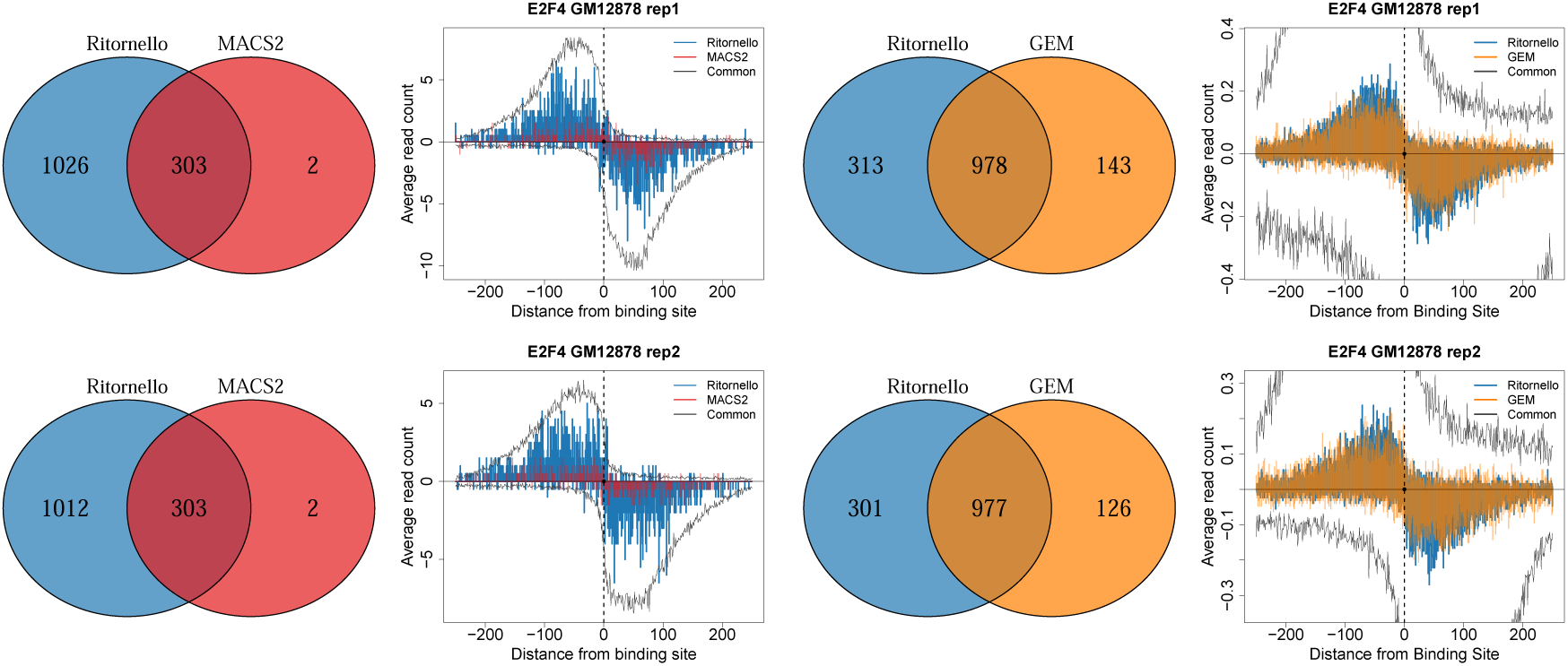
Pileup of read start positions for E2F4 reproducible peaks in GM12878 obtained by Ritornello only, by MACS2 only and by GEM only, where peaks within 100 bp of read length artifacts identified by Ritornello have been removed. The pileups of peaks common to Ritornello and MACS2 (left) or Ritornello and GEM (right) are shown in black.

**Figure S6:**
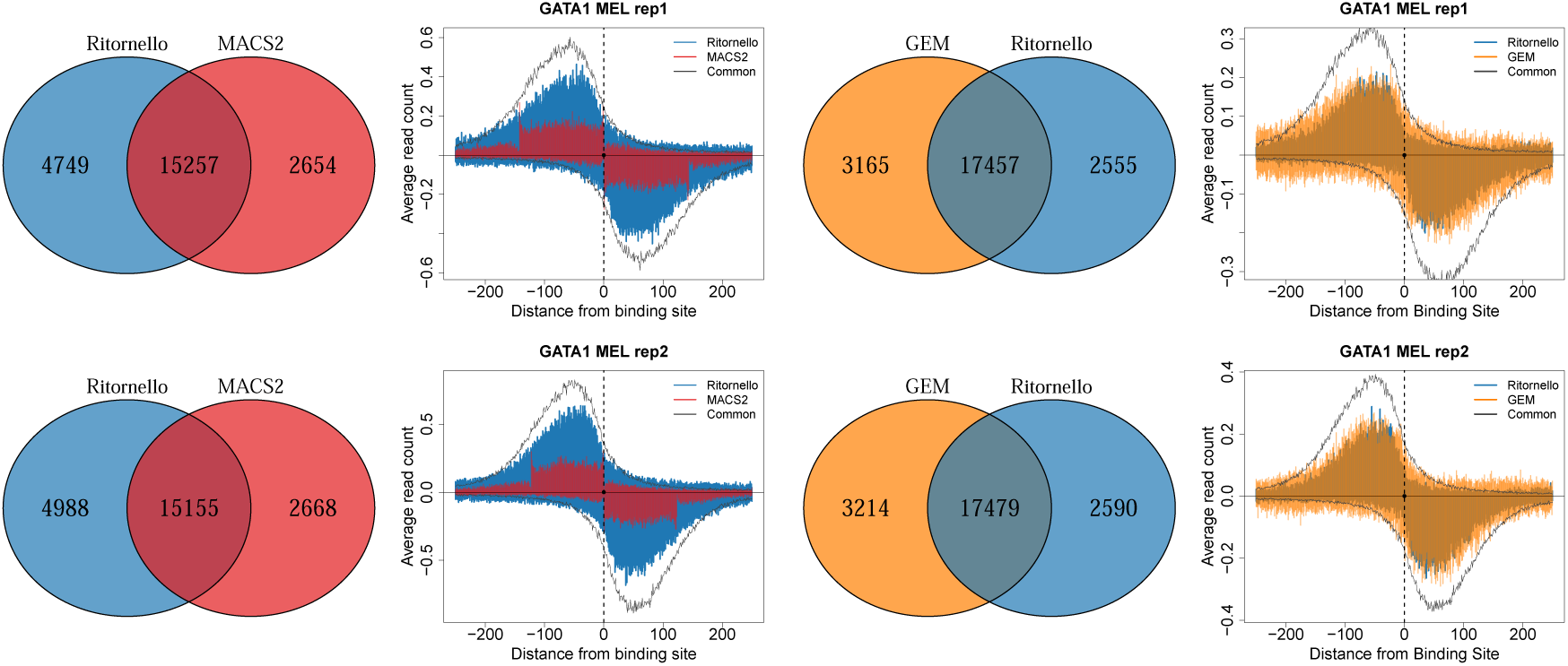
Pileup of read start positions for GATA1 reproducible peaks in MEL obtained by Ritornello only, by MACS2 only and by GEM only, where peaks within 100 bp of read length artifacts identified by Ritornello have been removed. The pileups of peaks common to Ritornello and MACS2 (left) or Ritornello and GEM (right) are shown in black. 4282 1608 995 Ritornello MACS2 −200

**Figure S7:**
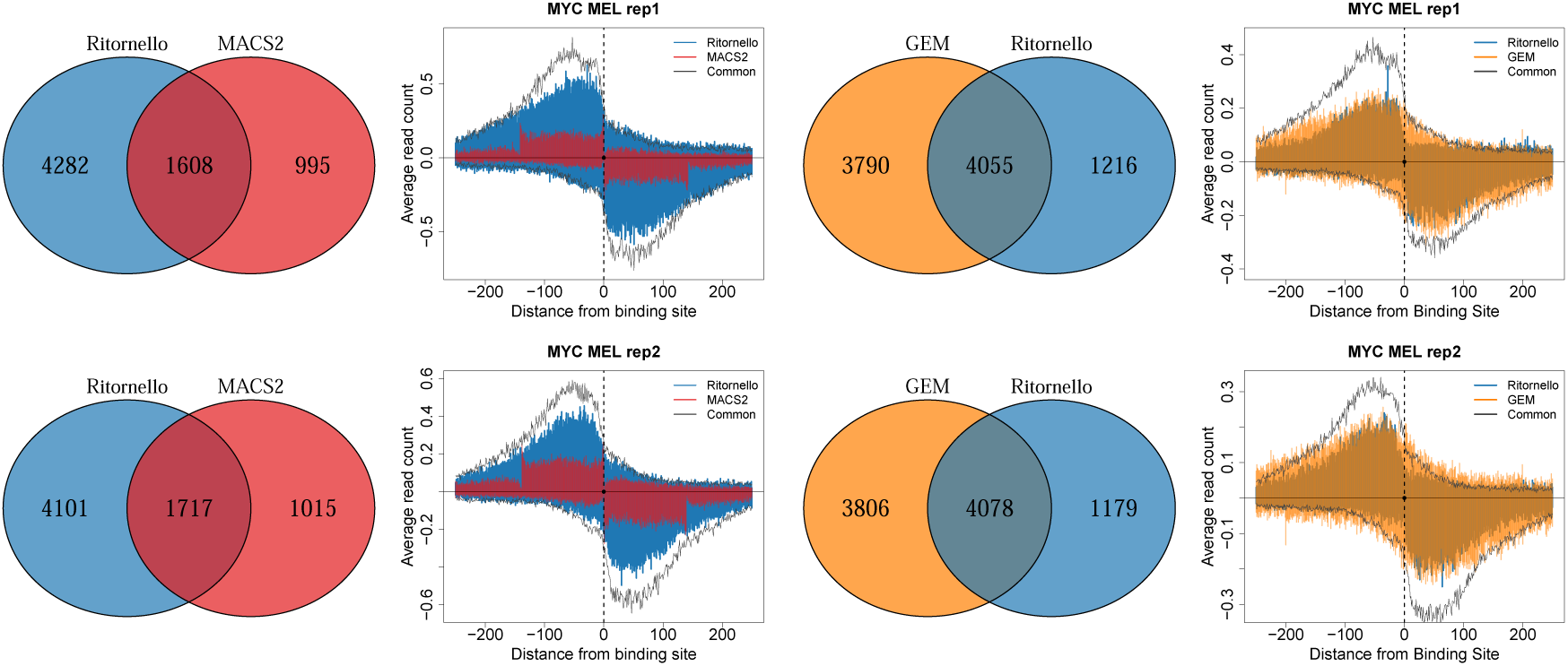
Pileup of read start positions for MYC reproducible peaks in MEL obtained by Ritornello only, by MACS2 only and by GEM only, where peaks within 100 bp of read length artifacts identified by Ritornello have been removed. The pileups of peaks common to Ritornello and MACS2 (left) or Ritornello and GEM (right) are shown in black.

**Figure S8:**
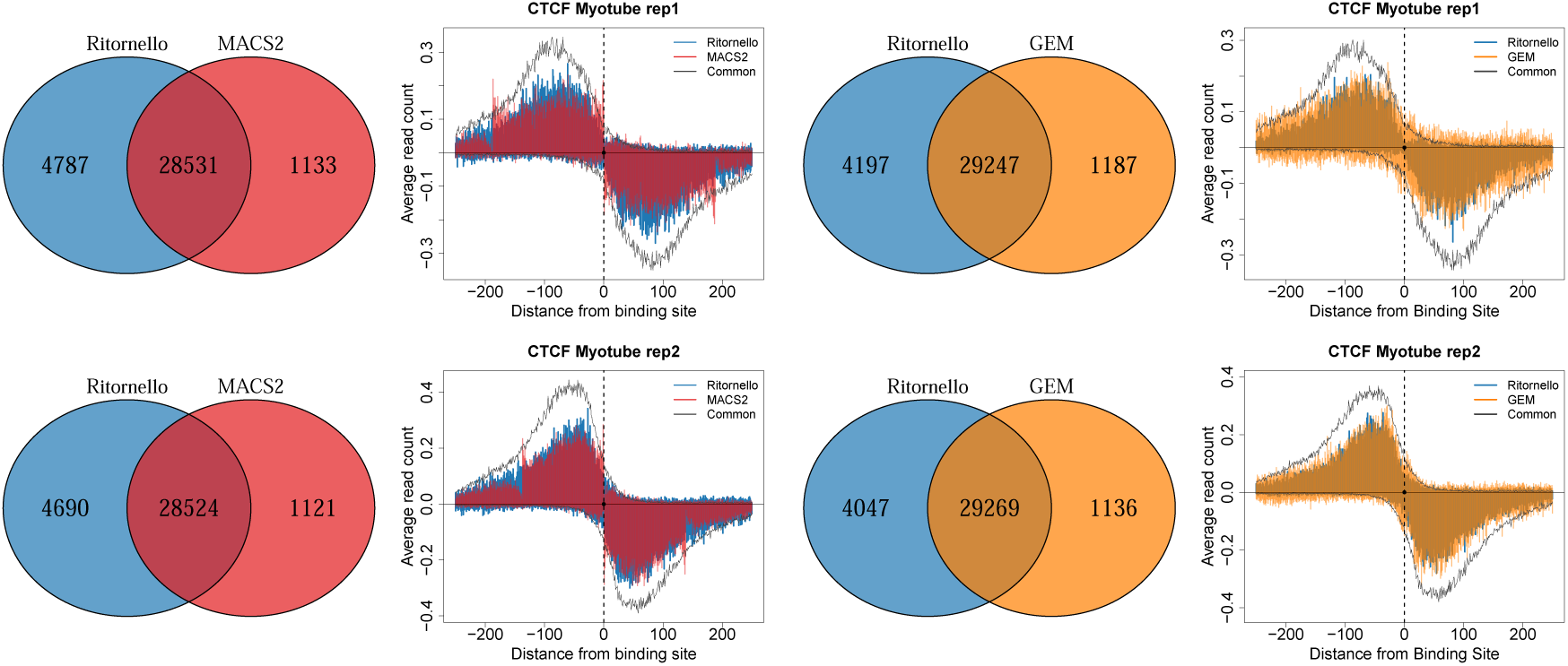
Pileup of read start positions for CTCF reproducible peaks in Myotube obtained by Ritornello only, by MACS2 only and by GEM only, where peaks within 100 bp of read length artifacts identified by Ritornello have been removed. The pileups of peaks common to Ritornello and MACS2 (left) or Ritornello and GEM (right) are shown in black.

**Figure S9:**
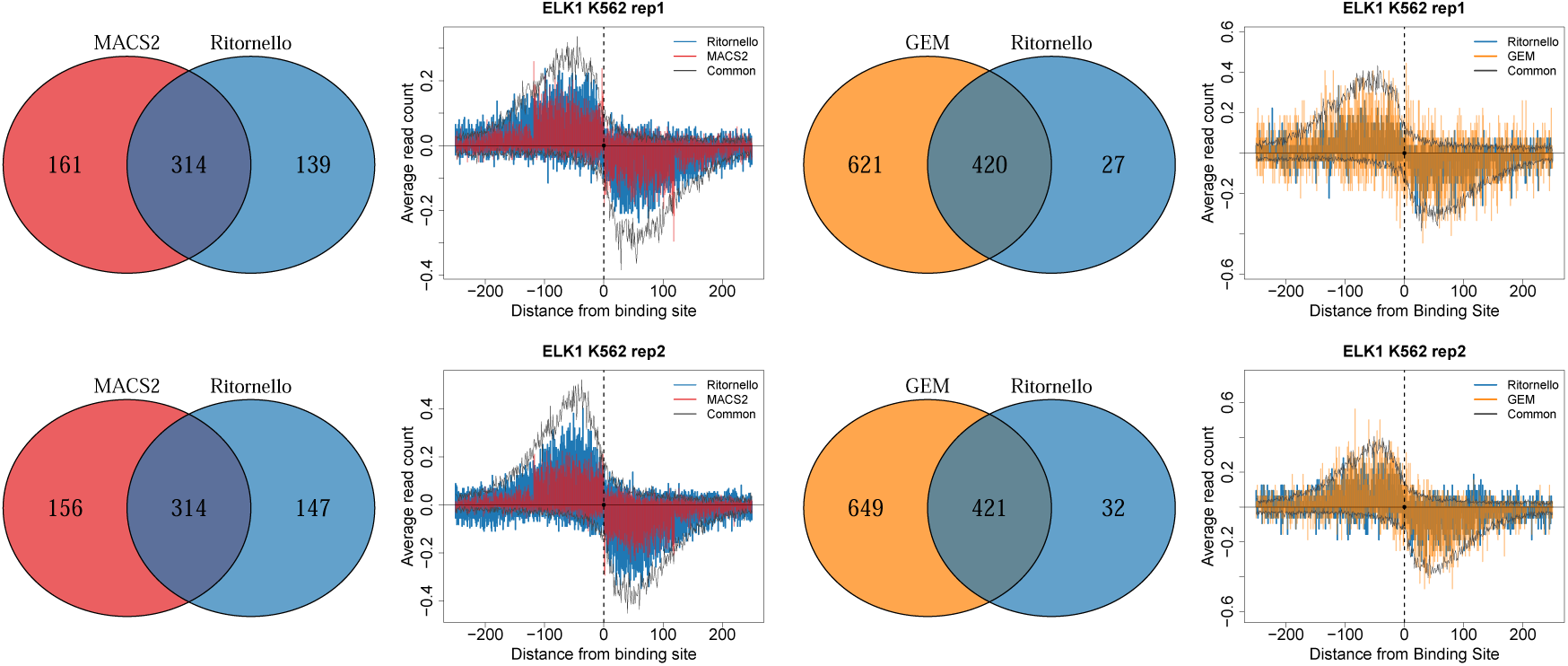
Pileup of read start positions for ELK1 reproducible peaks in K562 obtained by Ritornello only, by MACS2 only and by GEM only, where peaks within 100 bp of read length artifacts identified by Ritornello have been removed. The pileups of peaks common to Ritornello and MACS2 (left) or Ritornello and GEM (right) are shown in black.

**Figure S10:**
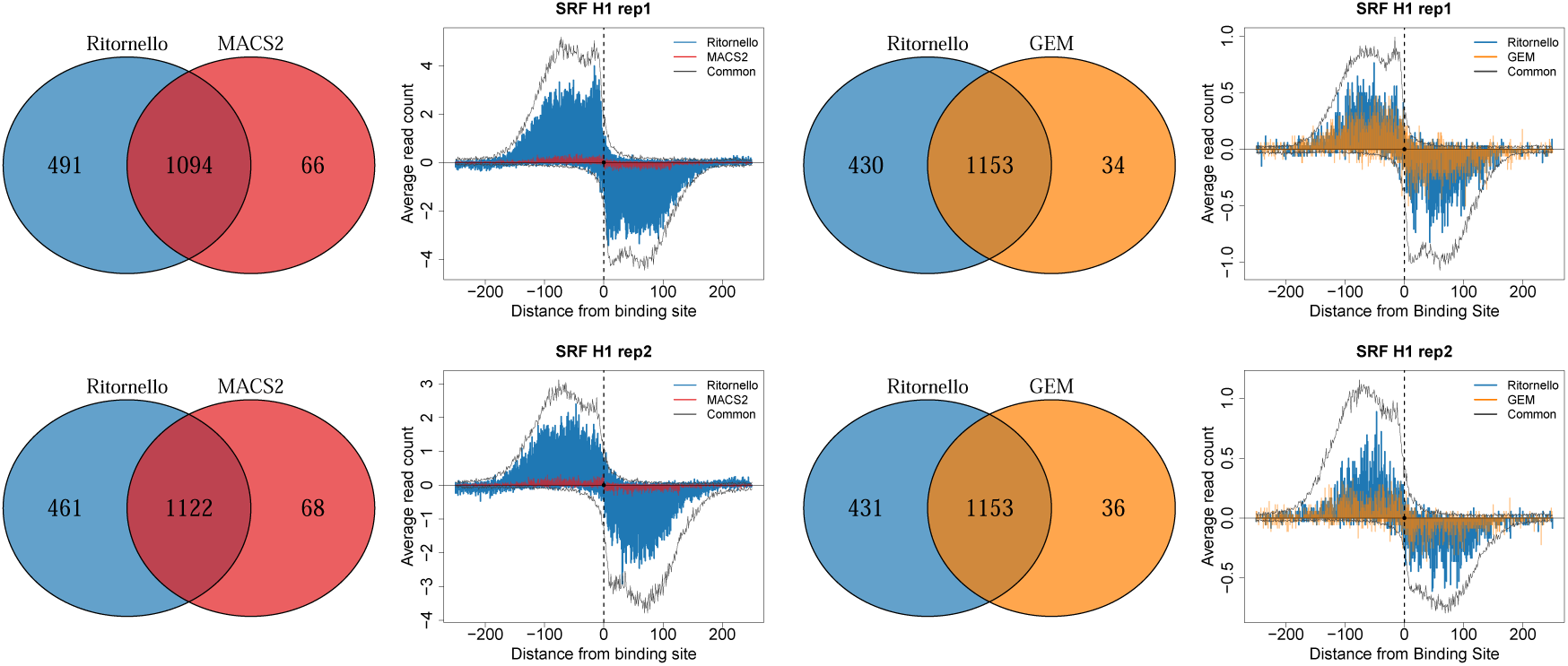
Pileup of read start positions for SRF reproducible peaks in H1 obtained by Ritornello only, by MACS2 only and by GEM only, where peaks within 100 bp of read length artifacts identified by Ritornello have been removed. The pileups of peaks common to Ritornello and MACS2 (left) or Ritornello and GEM (right) are shown in black.

**Figure S11:**
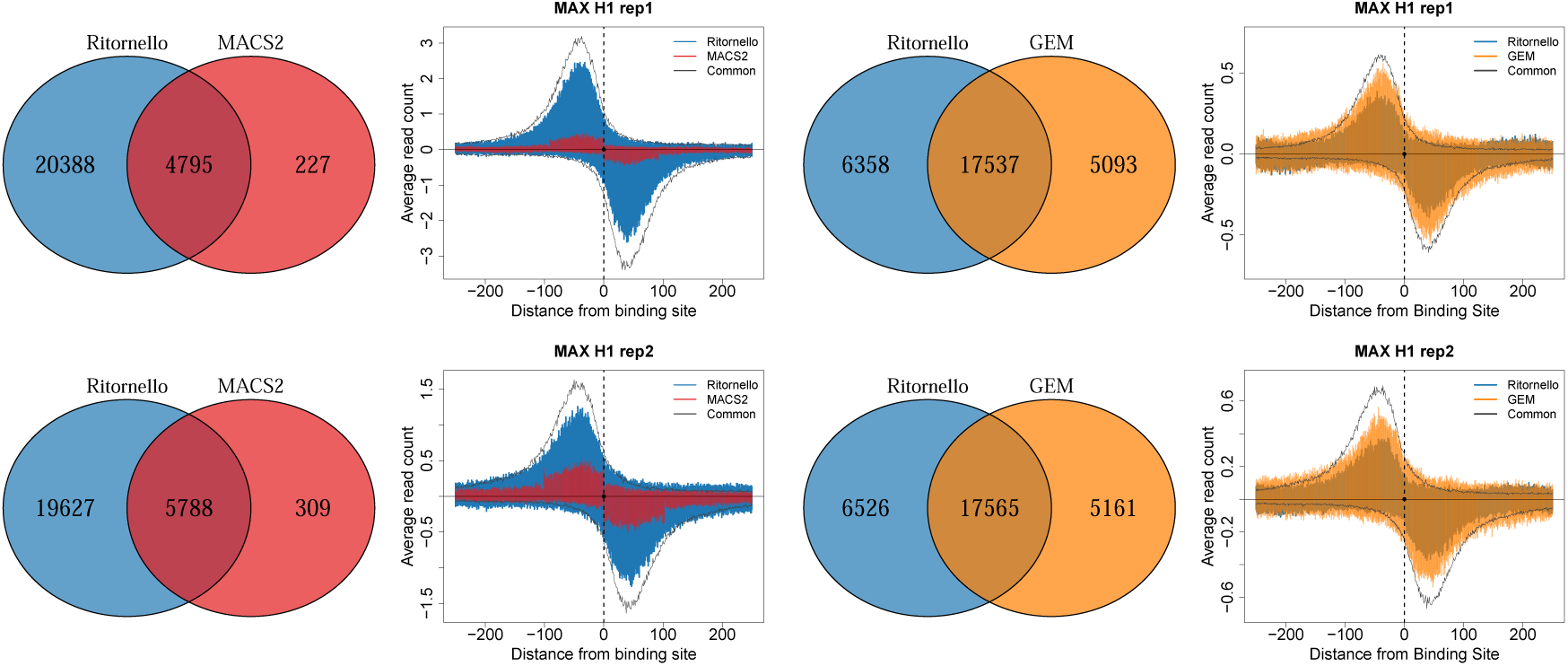
Pileup of read start positions for MAX reproducible peaks in H1 obtained by Ritornello only, by MACS2 only and by GEM only, where peaks within 100 bp of read length artifacts identified by Ritornello have been removed. The pileups of peaks common to Ritornello and MACS2 (left) or Ritornello and GEM (right) are shown in black.

**Figure S12:**
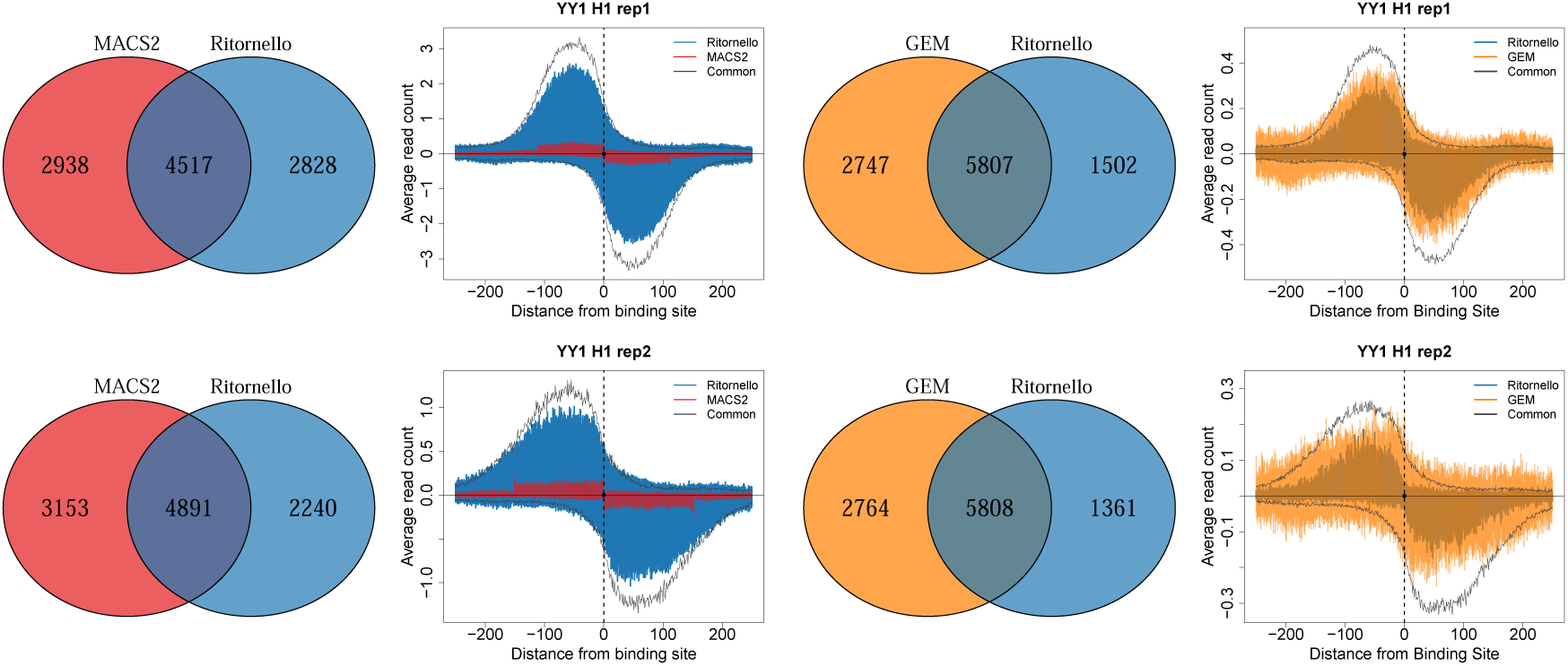
Pileup of read start positions for YY1 reproducible peaks in H1 obtained by Ritornello only, by MACS2 only and by GEM only, where peaks within 100 bp of read length artifacts identified by Ritornello have been removed. The pileups of peaks common to Ritornello and MACS2 (left) or Ritornello and GEM (right) are shown in black.

**Figure S13:**
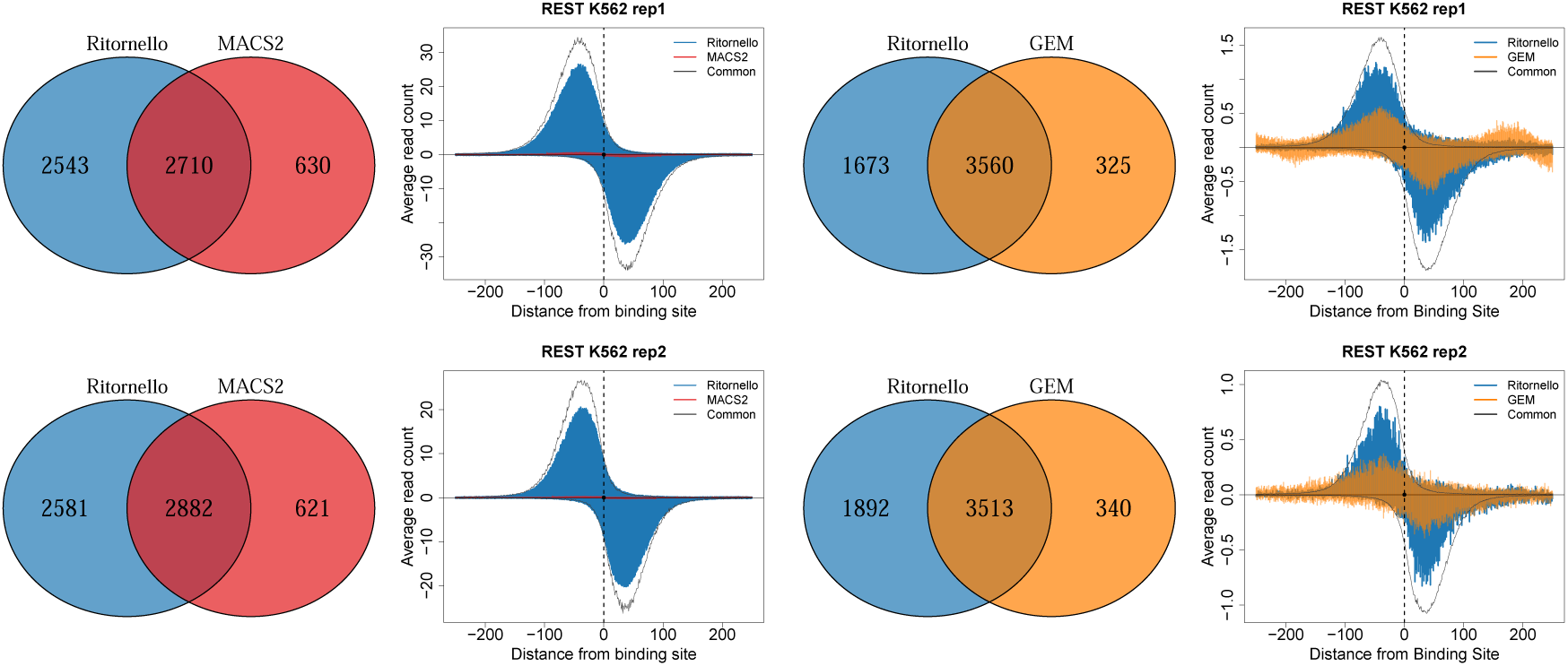
Pileup of read start positions for REST reproducible peaks in K562 obtained by Ritornello only, by MACS2 only and by GEM only, where peaks within 100 bp of read length artifacts identified by Ritornello have been removed. The pileups of peaks common to Ritornello and MACS2 (left) or Ritornello and GEM (right) are shown in black.

